# Clinically-driven design of synthetic gene regulatory programs in human cells

**DOI:** 10.1101/2021.02.22.432371

**Authors:** Divya V. Israni, Hui-Shan Li, Keith A. Gagnon, Jeffry D. Sander, Kole T. Roybal, J. Keith Joung, Wilson W. Wong, Ahmad S. Khalil

## Abstract

Synthetic biology seeks to enable the rational design of regulatory molecules and circuits to reprogram cellular behavior. The application of this approach to human cells could lead to powerful gene and cell-based therapies that provide transformative ways to combat complex diseases. To date, however, synthetic genetic circuits are challenging to implement in clinically-relevant cell types and their components often present translational incompatibilities, greatly limiting the feasibility, efficacy and safety of this approach. Here, using a clinically-driven design process, we developed a toolkit of programmable synthetic transcription regulators that feature a compact human protein-based design, enable precise genome-orthogonal regulation, and can be modulated by FDA-approved small molecules. We demonstrate the toolkit by engineering therapeutic human immune cells with genetic programs that enable titratable production of immunotherapeutics, drug-regulated control of tumor killing *in vivo* and in 3D spheroid models, and the first multi-channel synthetic switch for independent control of immunotherapeutic genes. Our work establishes a powerful platform for engineering custom gene expression programs in mammalian cells with the potential to accelerate clinical translation of synthetic systems.

## Main

The recent emergence of gene and cell-based therapies is being driven by their potential to enact precise and sophisticated responses to disease (*1*, *2*). One powerful example is chimeric antigen receptor (CAR) T cell immunotherapy, in which T cells are redirected to attack tumors by genetically engineering them to express artificial antigen-targeting receptors. This therapeutic paradigm is beginning to show clinical promise in treating certain cancers, leading to several approved cancer therapies (*3*). Yet while gene and cell-based therapies in general have tremendous potential to combat complex diseases, their impact has been limited by our inability to safely, effectively, and predictably control therapeutic cellular functions with engineered genetic systems. For example, engineered T cells also display adverse, sometimes fatal side effects due to issues related to off-target toxicity and over-activation (*3*–*5*), and have limited clinical efficacy for most solid tumors (*6*). Such limitations have motivated recent efforts in mammalian synthetic biology to design synthetic genetic programs that confer human cells with new capabilities and enable precise, context-specific control over therapeutic functions (*7*–*11*). While such examples represent significant advances, they also highlight fundamental challenges to realizing the compelling vision of implementing clinically-viable, therapeutic circuits in mammalian cells and eventually in humans (*2*, *12*): even simple synthetic circuits are difficult to implement in primary human cells and, critically, most are currently designed with regulatory components that present fundamental clinical incompatibilities. Overall, we lack versatile and clinically-suitable synthetic toolkits with which to reliably engineer relevant human cell types and fully unlock their therapeutic potential.

Perhaps the most established and effective method for implementing synthetic circuits to control mammalian cell behavior is using transcription regulation. Decades of research have illuminated design principles of transcription regulation (*13*), and established ways to engineer new conditions under which genes are expressed. These efforts have led to the development of a small, specialized, and widely-used set of artificial regulators based on microbial-derived transcription factors (TetR, Gal4) and viral activators (VP16, VP64), which exhibit robust functionality across many cell types and, in the case of TetR/tTa, can be induced by a small molecule antibiotic (*14*, *15*). However, the limited number of these regulators drastically restricts circuit linkages and the number of therapeutic genes that can be controlled, and they are challenging to reprogram for new regulatory specificities. Critically, their non-mammalian origins present clinical hurdles for therapies that depend on persistent expression: expression of TetR/tTa, for example, triggers immune responses in nonhuman primates (*16*, *17*). As such, though widely-used for decades, there are no clinically-approved genetic switches based on TetR/tTa. The emergence of programmable DNA-targeting elements, in particular the bacterial CRISPR/Cas9 system, has provided new methods for gene expression modulation and synthetic circuit design (*18*–*24*). However, the large genetic size of Cas9 poses a constraint on what can be designed and delivered to primary human cells, and its high immunogenic potential is well-documented (*25*–*28*). Thus, it would be powerful to establish a synthetic toolkit from first-principles that enables the development and translation of custom gene expression programs for clinically-relevant engineering contexts.

We outlined four basic principles for clinically-driven design of synthetic regulatory programs (**Fig. 1A**): (1) **Human-based** – prioritizing human-derived proteins when possible to minimize immunogenic potential. While some specialized human-based tools have been developed (*29*), they lack the scalability and programmability required of a foundational transcriptional toolkit for cell engineering. (2) **Orthogonal** – components with programmable, unique specificities that minimize cross-talk with native regulation. (3) **Regulatable** – systems that can be controlled with safe, clinically-suitable small molecules (and intrinsic biotic signals). (4) **Compact** – minimized genetic footprints for efficient delivery into primary human cells and tissues. Guided by these principles, we sought to develop – from primary sequence – a toolkit of compact, human-based synthetic transcription regulators that could readily enable engineering of custom gene expression circuits and programs in clinically-relevant human cells. As a building block for human-based regulators, we focused on Cys2His2 zinc fingers (ZFs), which provide the best balance of clinical favorability and programmability among known DNA-targeting elements. ZFs are small (~30 amino acid) domains that bind to ~3 bps of DNA (*30*, *31*). They are the most prevalent DNA-binding domain (DBD) found in human transcription factors (TFs) (*32*), suggesting they represent a highly flexible solution to DNA recognition with low immunogenicity potential. Indeed, a first-generation artificial ZF-based regulatory system showed multi-year functionality in non-human primates with no apparent immunogenicity (*33*). Moreover, individual ZF domains can be reprogrammed to recognize new motifs, and concatenated to generate proteins capable of specifically targeting longer DNA sequences (*34*–*37*). While these domains have been applied to generate tools for targeted endogenous modification and manipulation (*38*–*45*), ZFs have yet to be fashioned into a set of orthogonal and composable synthetic regulators with specificities optimized for the human genome. We sought to develop a protein engineering workflow that could fill this important gap and generate a versatile synthetic toolkit that satisfies the above criteria (**Fig. S1A**).

**Fig. 1.**
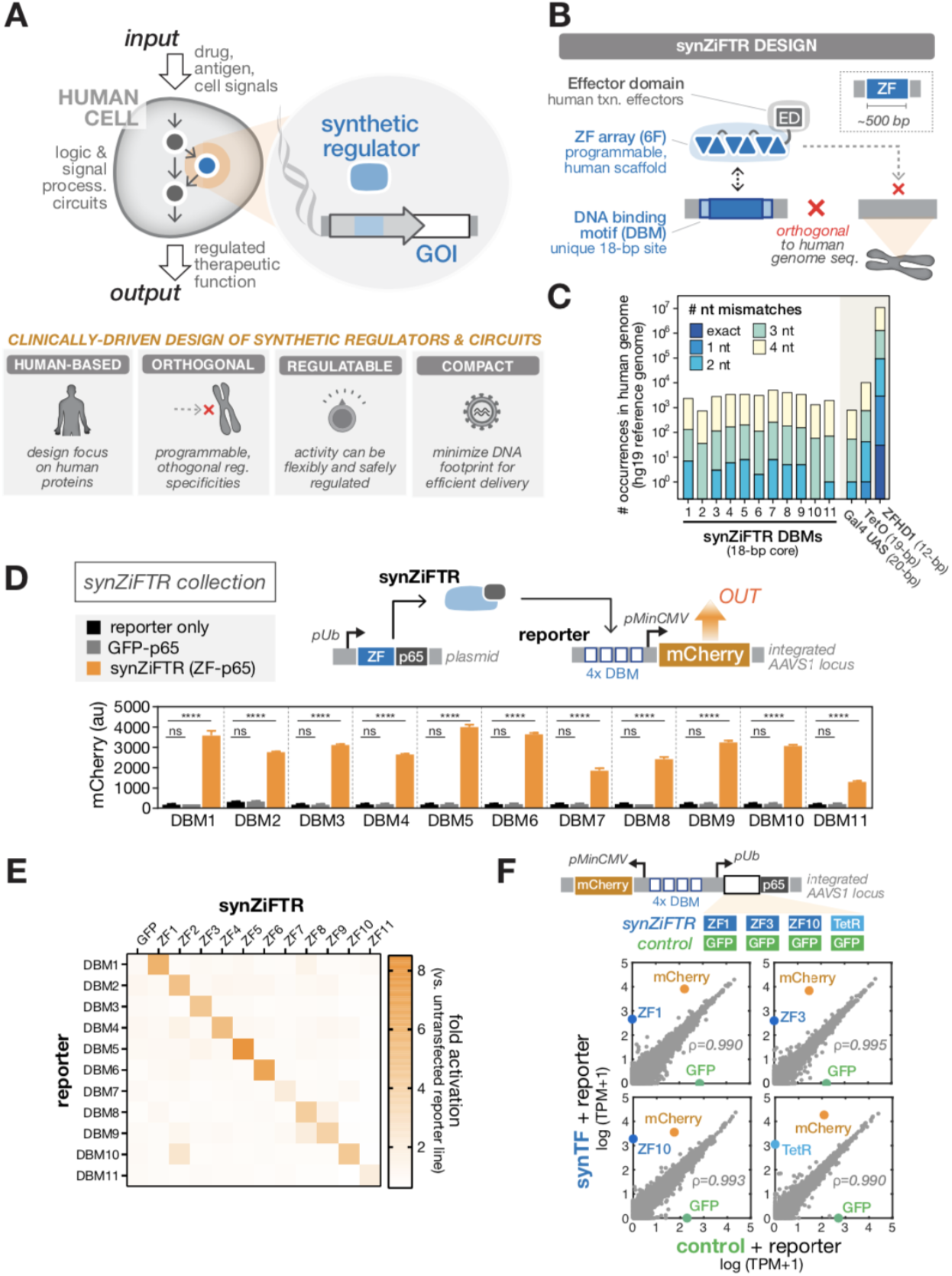
Clinically-driven design of compact, human, synthetic transcription regulators (synZiFTRs) (**A**) A goal of mammalian synthetic biology is to implement synthetic regulatory circuits that enable diverse forms of control over therapeutic functions in human cells (left). Principles of clinically-driven synthetic design (right). (**B**) Design of synZiFTRs. SynZiFTRs have a modular design based on compact, human-derived protein domains. An engineered ZF array mediates interactions with a unique, human genome-orthogonal DNA-binding motif (DBM), and human-derived effector domains (EDs) are used to locally control transcription functions. (**C**) Prevalence of synZiFTR recognition elements in the human genome. Occurrences of exact and increasingly mismatched sequences for each synZiFTR DBM and response elements from common artificial regulators (Gal4 UAS, TetO, ZFHD1). (**D**) SynZiFTRs strongly activate gene expression at corresponding response promoters. Response element vectors were stably-integrated into HEK293FT cells to generate reporter lines for each synZiFTR (ZF-p65 fusion). SynZiFTR (or control) expression vectors were transfected into corresponding reporter lines, and mCherry was measured by flow cytometry after 2 days. Bars represent mean values for three measurements ± SD. Statistics represent one-way ANOVA with Dunnett’s Multiple Comparisons; ns: not significant; ****: p < 0.0001. pUb, Ubiquitin C promoter; pMinCMV, minimal CMV promoter; p65, aa361-551. (**E**) SynZiFTRs have mutually orthogonal regulatory specificities. Each synZiFTR expression vector was transfected into every reporter line, and mCherry was measured by flow cytometry after 2 days. Fold activation levels represent mean values for three biological replicates. (**F**) SynZiFTRs have highly specific regulatory profiles in human cells. Correlation of transcriptomes from RNA-sequencing measurements of HEK293FT cells stably expressing synZiFTR or TetR-p65 versus a GFP-p65 control. Points represent individual transcript levels normalized to TPM, transcripts per kilobase million, averaged between two technical replicates. Pearson correlation coefficient was calculated for native (grey) transcripts. See **fig. S3** for extended analyses.

To design genome-orthogonal regulators, we leveraged an archive of engineered two-finger (2F) units, based on the canonical human Egr1 scaffold. These units were pre-generated using selection-based methods and explicitly account for context-dependent effects between adjacent fingers (*35*, *37*). By linking 2F units (each recognizing 6-bp subsites) using flexible ‘disrupted’ linkers (*46*), it is possible to construct functional six-finger (6F) arrays capable of recognizing 18 bps, a length for which a random sequence has a high probability of being unique in the human genome (**Fig. 1B, S1B**). We prioritized 6-bp subsites that are underrepresented in the human genome and selected arrays to minimize identity with the human genome; this yielded 11 targetable synthetic DNA-binding motifs (DBMs) (**Fig. 1C, S1C-D, Methods**). We next sought to engineer compact, human-based, <u>syn</u>thetic <u>Zi</u>nc <u>F</u>inger <u>T</u>ranscription <u>R</u>egulators (**synZiFTRs**) capable of strong and specific regulation at these synthetic *cis*-elements. We fused ZFs predicted to bind each DBM to the human p65 activation domain and screened for the most active synZiFTR candidate for each DBM in HEK293FT reporter lines (**Fig. S2A-C**). Our selected synZiFTRs strongly activate corresponding, but not non-cognate, reporters (**Fig. 1D-E**). Finally, to evaluate the impact of our synthetic regulators on native regulation, we performed RNA-sequencing analysis on cell lines expressing three representative synZiFTRs (ZF1, ZF3, ZF10) and benchmarked these against a TetR-based activator. SynZiFTR regulation profiles are highly specific, minimally affecting native transcript profiles and importantly compare favorably with that of TetR (**Fig. 1F, S3**). These results establish a collection of compact, human-based, and genome-orthogonal synZiFTRs optimized for artificial gene expression control in human cells.

Regulatory systems that offer precise control over the timing, level, and context over which therapeutic genes are expressed could enable safer and more effective gene and cell-based therapies (*2*). One promising approach is exogenous gene expression control using small molecules, which could be administered systemically or locally to switch ON and allow titratable control over therapeutic gene products. To date, there are no clinically-approved small molecule-regulated genetic switches, in part because systems that have been developed utilize toxic or pharmacodynamically-unfavorable small molecules. We focused on developing small molecule-inducible synZiFTRs around compounds that are clinically-approved or otherwise known to have favorable safety profiles (**Fig. 2A**). We selected three classes of small molecules, which regulate protein activity through distinct mechanisms, offering the potential for up to three orthogonal channels of gene expression control (**Fig. 2B, S4A**): 1) Grazoprevir (GZV), an FDA-approved antiviral drug from a family of protease-inhibiting compounds, which has an exceptional safety profile and is commonly taken at a high dose (100 mg/day) for up to 12 weeks (*47*). Addition of GZV stabilizes synZiFTRs incorporating the NS3 self-cleaving protease domain (from hepatitis C virus (HCV)), driving gene transcription (*48*, *49*). 2) 4-Hydroxytamoxifen (4OHT), a metabolite of the widely prescribed breast cancer drug tamoxifen that selectively modulates the nuclear availability of molecules fused to estrogen receptor variants, such as ERT2 (*50*, *51*). 3) Abscisic acid (ABA), a plant hormone naturally present in many plant-based foods and classified as non-toxic to humans, which mediates conditional binding of the domains ABI and PYL to reconstitute an active synZiFTR (*52*).

**Fig. 2.**
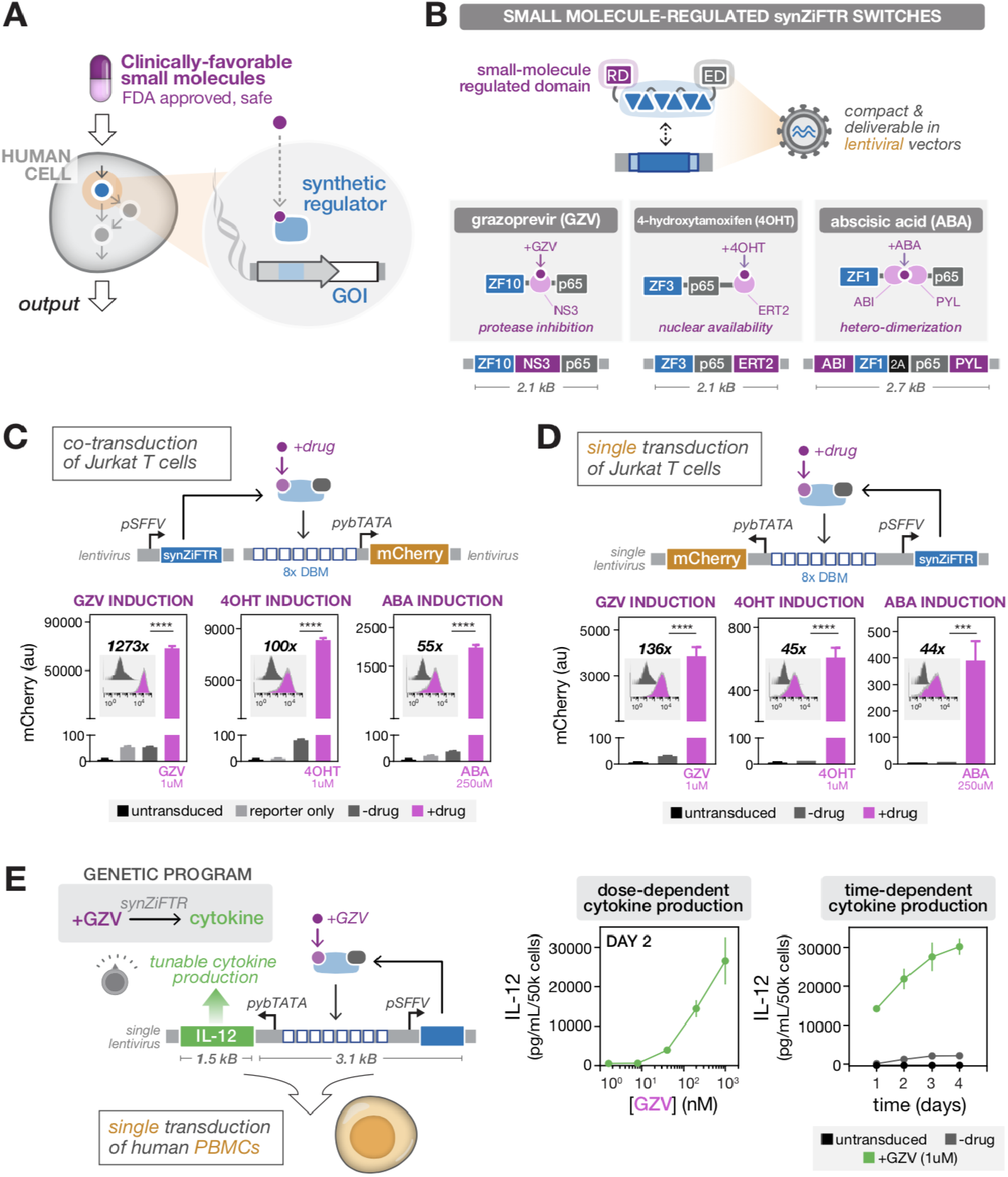
SynZiFTR switches controlled by clinically-favorable compounds enable inducible and titratable control of gene expression in human immune cells. (**A**) Exogenous control of therapeutic genetic programs can be enabled by synZiFTRs that respond to clinically-favorable pharmaceutical compounds. (**B**) Design of three classes of synZiFTR switches controlled by orthogonal small molecules: grazoprevir (GZV), 4-hydroxytamoxifen (4OHT), abscisic acid (ABA). NS3, hepatitis C virus NS3 protease domain; ERT2, human estrogen receptor T2 mutant domain; ABI, ABA-insensitive 1 domain (aa 126-423); PYL, PYR1-like 1 domain (aa 33-209); 2A, 2A self-cleaving peptide. (**C**) Optimized synZiFTR switches enable inducible gene expression control in Jurkat T cells with strong activation profiles. Jurkat T cells were co-transduced with reporter and synZiFTR expression lentiviral vectors in an equal ratio. mCherry was measured by flow cytometry 4 days following induction by small molecules at indicated concentrations. Bars represent mean values for three measurements ± SD. Statistics represent two-tailed Student’s t test; ***: p < 0.001; ****: p < 0.0001. Histograms show absolute levels and mean fold activation for one representative measurement (insets). pSFFV, Spleen Focus-Forming Virus promoter; pybTATA, synthetic YB_TATA promoter. (**D**) Optimized synZiFTR switches enable design of compact, single lentiviral vectors for strong and inducible gene expression control in Jurkat T cells. (**E**) Development of a compact, GZV-regulated synZiFTR program for tight, titratable control of IL-12 cytokine production in primary human immune cells. Human primary peripheral blood mononuclear cells (PBMCs) were activated and transduced with a single lentiviral vector encoding single-chain IL-12 payload and synZiFTR expression cassettes (see **Methods**). IL-12 production was measured by ELISA at specified time points following induction (with or without 1 uM GZV). Points represent mean values for three measurements ± SD.

To evaluate the ability of these small molecules to control synZiFTR activity, we constructed GZV-, 4OHT- and ABA-inducible switches based on distinct ZFs (ZF1, ZF3, ZF10), and encoded these and associated reporters on lentiviral vectors (**Fig. S2B, S4A**). We then performed a series of characterization studies by co-transducing Jurkat T cell lines with synZiFTR and reporter constructs (**Fig. S4B**). All three systems exhibited titratable control of reporter output, minimal leakage relative to reporter-only cells, strong dynamic ranges, and returned to basal (OFF) levels upon removal of inducer (**Fig. S4C**); furthermore, the small molecules only activated respective synZiFTRs (**Fig. S4D**). Since the small molecules function through distinct and orthogonal regulatory mechanisms, we realized that they could be multiplexed within the same synZiFTR molecule to enable more complex forms of temporal gene expression control, including ON/OFF switching. For example, we built an ON/OFF switch by incorporating the SMASh domain, a variant of NS3 that functions as a degron in the presence of GZV (*53*), onto an ERT2-synZiFTR (**Fig. S5A,B**). The switch is turned ON in the presence of 4OHT, and addition of GZV turns gene expression OFF, returning the system to basal levels rapidly (**Fig. S5C,D**). We also demonstrated a second type of ON/OFF switch, using 4OHT and GZV to respectively activate and degrade a synZiFTR repressor containing either human-derived KRAB or HP1α domains (**Fig. S5E,G**). Taken together, these results demonstrate that synZiFTR activity can be regulated by three, orthogonal, and clinically-favorable small molecules, which can be multiplexed to yield versatile genetic switch designs.

Before translating our synZiFTR systems to clinically-relevant cell types, we optimized response elements by screening arrangements of DBM arrays and minimal promoters to identify combinations that reduced basal expression and improved dynamic range (**Fig. S6**). Collectively, these studies led to an optimized design of GZV-, 4OHT-, and ABA-inducible synZiFTR systems that can be compactly encoded on either dual or single lentiviral vectors to enable strong, tight induction of gene expression (**Fig. 2C,D**). We then sought to determine whether synZiFTRs could be used to control expression of therapeutically-relevant payloads. For these studies, we chose to focus on engineering primary human immune cells, which have immense therapeutic potential but for which precise gene expression control remains highly challenging; thus, establishing our systems in these cells will provide a blueprint for translation into other challenging, clinically-relevant cell types. As an initial proof-of-principle, we selected the immunomodulatory factor, IL-12. IL-12 has potent anti-tumor activity, but overexpression of IL-12 can cause severe and fatal side effects because of dose-limiting toxicity, making it a promising candidate for synZiFTR-regulated expression control in engineered immune cells (*54*, *55*). We constructed a single-lentiviral vector encoding a GZV-regulated IL-12, which we used to transduce primary human immune cells (**Methods**). Significantly, our synZiFTR system enabled titratable control over IL-12 production in a GZV dose- and time-dependent manner (**Fig. 2E**). These results provide a promising basis for compact synZiFTR-based switches and programs capable of controlling therapeutically-relevant payloads.

Do synZiFTR programs drive clinically-relevant outputs? To answer this, we turned to the CAR T cell therapy paradigm, initially choosing to develop a synZiFTR-controlled anti-Her2 CAR (**Fig. 3A**). Her2 is a tyrosine kinase receptor that is overexpressed in many tumors, including a small subset of leukemia (*56*–*59*). We have previously successfully used this anti-Her2 CAR in a xenograft liquid tumor model, thus providing a convenient platform to evaluate the efficacy of our synZiFTR circuits (*60*). We generated a GZV-inducible anti-Her2 CAR system, which enabled inducer-dependent CAR expression in primary human T cells, notably to levels comparable to a standard constitutively-expressed CAR and with minimal output in the absence of inducer (**Fig. 3B**). When co-cultured with a Her2-overexpressing (HER2+) NALM6 leukemia cell line (**Fig. S7C**), we found that synZiFTR-controlled CAR cells were capable of GZV-dependent activation and efficient tumor cell killing *in vitro* (**Fig. 3C, S8A**). Importantly, these synZiFTR programs are easily reconfigurable. By swapping the anti-Her2 CAR with an anti-CD19 CAR, we could reproduce these *in vitro* activity results for a second payload, confirming the generalizability of our system (**Fig. S7**).

**Fig. 3.**
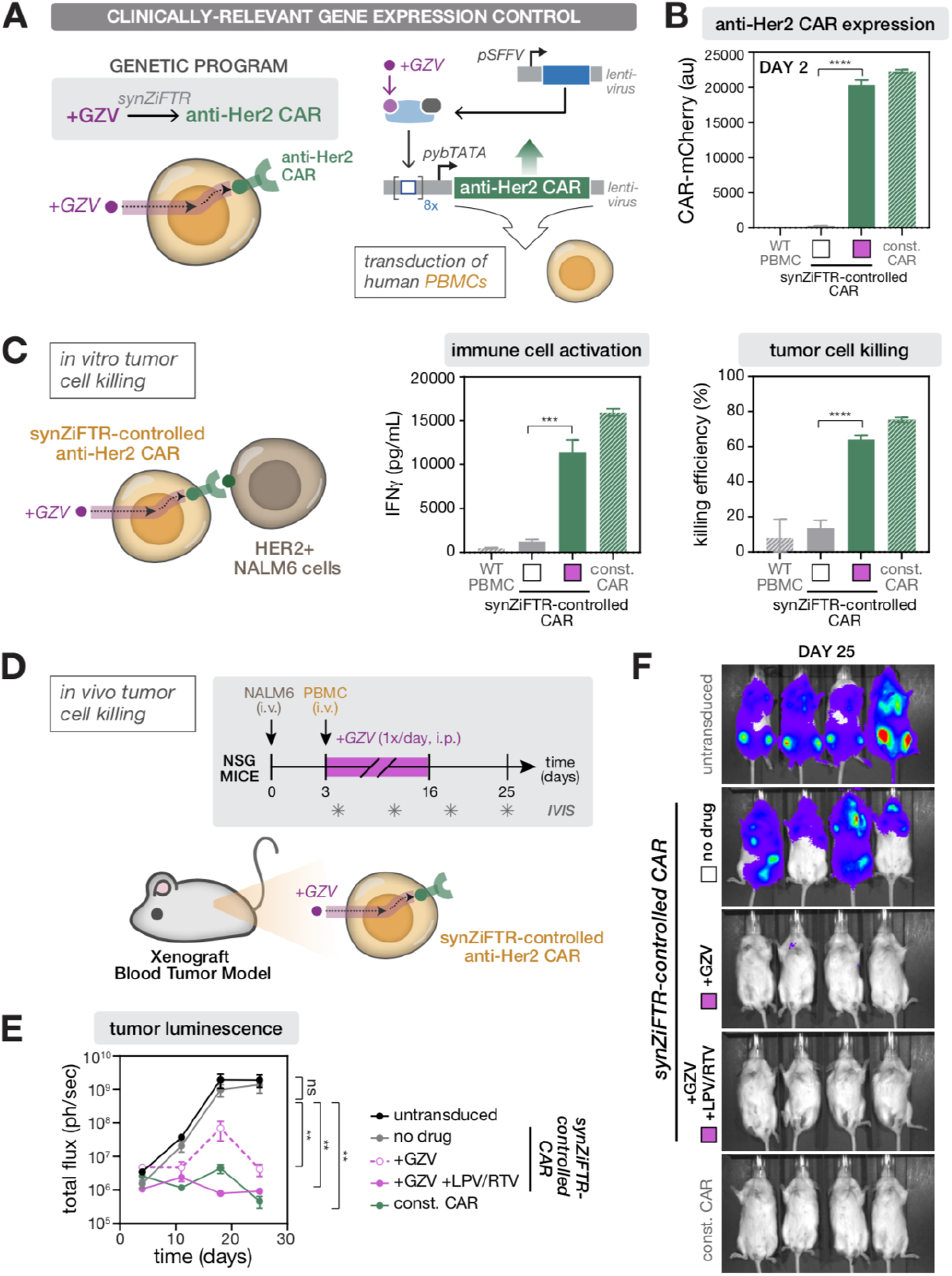
SynZiFTR programs enable inducible, clinically-relevant gene expression control over CAR T cell activity *in vivo*. (**A**) Conceptualization (left) and implementation (right) of a synZiFTR-controlled anti-Her2 CAR gene expression program. Human PBMCs were activated and co-transduced with equal ratios of lentiviral vectors encoding anti-Her2 CAR payload and synZiFTR expression cassettes (see **Methods**). (**B**) SynZiFTR program enables GZV-dependent CAR expression in primary human immune cells. Expression of anti-Her2 CAR-mCherry was measured by flow cytometry two days following induction (with or without 1 uM GZV). Const. CAR, constitutively expressed (pSFFV-CAR). Bars represent mean values for three measurements ± SD. Statistics represent two-tailed Student’s t test; ***: p < 0.001; ****: p < 0.0001. (**C**) SynZiFTR-CAR program enables GZV-dependent immune cell activation and tumor cell killing *in vitro*. SynZiFTR-controlled CAR cells (pre-induced with or without 1 uM GZV for 2 days) were co-cultured with HER2+ NALM6 target leukemia cells in a 1:1 ratio (left). IFNγ secretion from activated immune cells was measured by ELISA (center) and tumor cell killing by flow cytometry (right), one day following co-culturing. (**D**) Testing *in vivo* efficacy of synZiFTR-controlled CAR T cells using a xenograft tumor mouse model. Timeline of *in vivo* experiments, in which NSG mice were injected i.v. with luciferized HER2+ NALM6 cells to establish tumor xenografts, followed by treatment with PBMCs. GZV was formulated alone or in combination with LPV/RTV and administered i.p. daily over 14 days. Mice were imaged weekly on days 4, 11, 18, 25 to monitor tumor growth via luciferase activity. GZV, 25 mg/kg. LPV/RTV, 10 mg/kg. (**E**) Tumor burden over time, quantified as the total flux (photons/sec) from the luciferase activity of each mouse using IVIS imaging. Points represent mean values ± SEM (n=4 mice per condition). Statistics represent two-tailed, ratio paired Student’s t test; ns: not significant; **: p < 0.01. (**F**) IVIS imaging of mouse groups treated with (1) untransduced PBMCs, (2) synZiFTR-controlled CAR cells, (3) synZiFTR-controlled CAR cells with GZV, (4) synZiFTR-controlled CAR cells with GZV+LPV/RTV, (4) constitutive CAR cells. (n=4 mice per condition).

Finally, we tested the *in vivo* efficacy of synZiFTR-controlled CAR cells using a simple xenograft blood tumor model (*60*) (**Fig. 3D, Methods**). For mice receiving synZiFTR-controlled anti-Her2 CAR T cells, GZV was dosed every day at 25 mg/kg for 14 days, either alone or in combination with lopinavir/ritonavir (LPV/RTV, 10 mg/kg), an antiretroviral drug cocktail known to increase GZV bioavailability (*61*). Importantly, GZV inducer combinations have no effect on cells on their own (**Fig. S8A**), and measurements of mouse body weight over the course of the experiment confirmed that daily inducer injections were not toxic (**Fig. S7B**). Tumor growth was monitored via IVIS imaging of luciferase-expressing HER2+ NALM6 cells over the course of 25 days (**Fig. 3E**). Mice receiving synZiFTR-controlled CAR cells and treated with GZV or GZV+LPV/RTV were able to clear the tumor, while those not treated with inducer resulted in high tumor burdens (**Fig. 3E-F**). Interestingly, while both inducer conditions ultimately led to tumor eradication, clearance rates were faster with the cocktail, on par with the constitutive CAR positive control and consistent with the ability of LPV/RTV to increase GZV bioavailability (**Fig. 3E, S8C**). This suggests that leveraging pharmacokinetic considerations could provide yet another way to tune synZiFTR circuit output *in vivo*. Overall, these results establish genetic switches for inducible CAR expression, and demonstrate that synZiFTRs can be used to program drug-dependent control over T cell therapeutic activity *in vivo*.

There is an increasing recognition that “co-engineering” strategies will be crucial to enhancing the efficacy of gene and cell therapies. For example, co-engineering tumor-targeting immune cells with immunomodulatory factors that can combat immune suppression, promote tumor homing, and/or enhance immune responses will be key to improving the efficacy of CAR T cells against solid tumors, as well as to realizing the next-generation of CAR T cell therapies for other diseases, such as autoimmunity and heart disease (*62*–*64*). Pleiotropic factors, such as IL-12, have been explored for immunotherapy due to their anti-tumor properties. Other interesting candidates include IL-4, which may facilitate tissue repair and suppress inflammation (*65*). Importantly, these cytokines have diverse, contextual, and dose-dependent functions within the human body, once again motivating the critical need for precise spatiotemporal control of gene expression. Our synZiFTR platform provides a promising basis for next-generation coengineering strategies by enabling custom, “multi-channel” synthetic programs, in which therapeutic genes are independently regulated by orthogonal small molecules (**Fig. 4A**). As a proof-of-principle, we constructed a two-channel synthetic system in which one channel is dedicated to controlling a tumor-targeting factor (e.g. CAR) and a second channel to controlling a desired immunomodulatory payload (e.g. cytokines). Specifically, our design was composed of a GZV-inducible synZiFTR (ZF10) switch controlling anti-Her2 CAR and a 4OHT-inducible synZiFTR (ZF3) switch controlling either IL-12 or IL-4. Following transduction of primary human immune cells with our synthetic two-channel systems, we found that addition of GZV and 4OHT led to the production of CAR and cytokines, respectively, with minimal apparent output in the no drug condition (**Fig. 4B, S9A**).

**Fig. 4.**
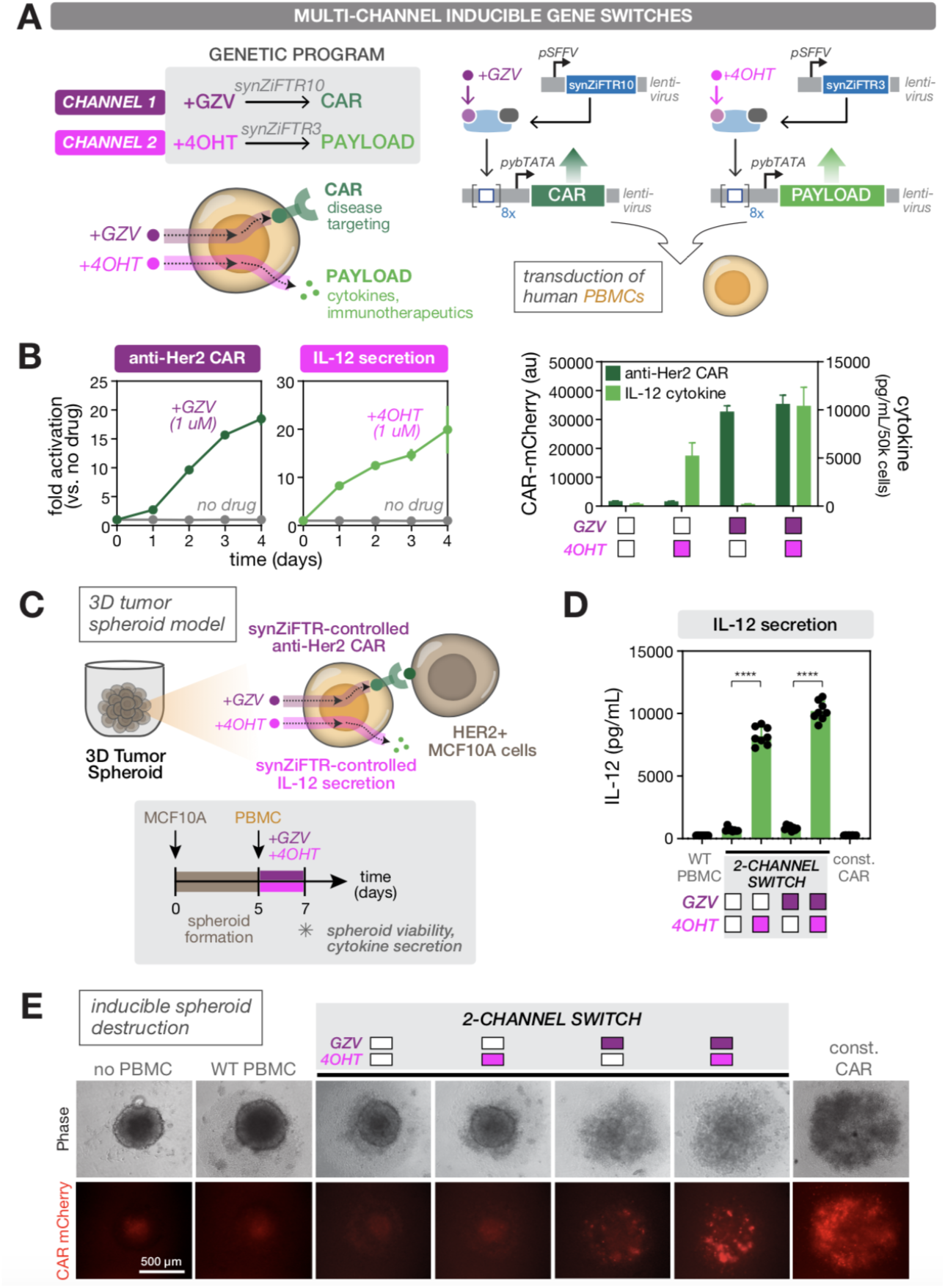
Development of a two-channel synthetic switch for independent, drug-regulated control of immunotherapeutic products. (**A**) Conceptualization (left) and implementation (right) of the two-channel synthetic system. Orthogonal small molecule-regulated synZiFTR switches (GZV-synZiFTR10 and 4OHT-synZiFTR3) are used to control expression of a CAR and an immunomodulatory payload, respectively. (**B**) Independent, drug-inducible control of anti-Her2 CAR and IL-12 in primary human immune cells. PBMCs were transduced with lentiviral vectors comprising the two-channel system, and induced with combinations of GZV and 4OHT. Anti-Her2-CAR-mCherry was measured by flow cytometry and cytokine secretion by ELISA. Points (left) and bars (right) represent mean values for three measurements ± SD. (**C**) Testing efficacy of two-channel synZiFTR switches using a 3D tumor spheroid model based on HER2+ MCF10A cells. Timeline of spheroid formation and experiments (bottom). +GZV, 1 uM; +4OHT, 1 uM. (**D**) 4OHT-inducible control over IL-12 secretion in 3D tumor spheroid co-culture. IL-12 was quantified by ELISA analysis of supernatant. Bars represent mean values ± SD (n=8 spheroids per condition). Statistics represent two-tailed Student’s t test; ****: p < 0.0001. (**E**) GZV-inducible control over spheroid destruction by engineered two-channel cells. Representative phase contrast and CAR-mCherry fluorescent images of spheroid morphology when co-cultured with two-channel inducible or control PBMCs. Clear disruption of the compact, rounded morphology is seen in conditions in which PBMC express the anti-HER2 CAR.

To demonstrate the efficacy of the two-channel switch in driving functional changes in cell behavior, we designed and developed a spheroid tumor target for the engineered immune cells. While spheroids are an imperfect model of *in vivo* solid tumors, they share notable morphological and behavioral similarities, including the development of oxygen and nutrient gradients, formation of a necrotic/apoptotic central core, and recapitulation of 3D cell-cell and cell-matrix interactions (*66*). As such, spheroids are thought to provide a more physiologic *in vitro* response to therapeutics than simple 2D monolayers, and are widely used to evaluate the efficacies of small molecule and cellular therapeutics in high-throughput (*67*). Here we designed and employed a 3D spheroid, based on HER2+ MCF10A breast mammary epithelial cells, to simultaneously probe CAR-mediated spheroid killing and immunomodulatory cytokine secretion (**Fig. 4C, S9, Methods**). We observed clear morphological differences between spheroids cocultured with CAR-expressing (+GZV or constitutive CAR) and non-expressing (-GZV and WT) cells. Specifically, while spheroids cultured with uninduced cells (-GZV) retained their compact rounded morphology, spheroids cultured with cells induced to express anti-HER2 CAR (+GZV) showed significant morphological disruption, including loss of their hallmark rounded shape and amorphous cell scattering throughout the well, both signs of spheroid fragmentation and disassembly that are indicative of CAR-mediated cell killing (**Fig. 4E**). These stark morphologic differences were observed over multiple biological replicates (**Fig. S9E**) and are consistent with our previous *in vitro* and *in vivo* tumor cell killing data (**Fig. 3C,E**). Supernatant analysis from the spheroid co-cultures showed significant IL-12 production only in 4OHT treated conditions, demonstrating 4OHT-mediated control over local production of the cytokine (**Fig. 4D**). Collectively, these results demonstrate spheroid killing and cytokine expression behaviors in the presence of GZV and 4OHT, respectively, illustrating the multi-channel control enabled by the synthetic switch. Importantly, inducers alone exhibited no morphological disruption indicative of spheroid killing (**Fig. S9D**). More broadly, these results demonstrate that synZiFTRs can be used to implement complex therapeutic gene expression programs in clinically-relevant cell types, and establish the first two-channel, synthetic switch for independent control of immunotherapeutic genes in primary human immune cells.

In this work, we outlined a set of minimal principles for clinically-driven design of synthetic regulatory programs in human cells. These principles were reflected in the choice of molecular building blocks and features that we prioritized in developing a versatile human synZiFTR toolkit. We based our synZiFTRs on engineered ZFs for their intrinsic flexibility in DNA targeting, compact size, human origins, and reported history of not eliciting unintended immune responses. Though not as easily programmable as CRISPR/Cas9 systems, ZFs have been shown to function in virtually every cell type in which they have been tested, their specificity can be reprogrammed, and they can be effectively used as building blocks for engineering highly specific and active regulators when workflows are developed that incorporate context-dependencies and other ZF design criteria. Here, we demonstrated the development of, to our knowledge, the first collection of human genome-orthogonal, ZF-based synthetic regulators. Other efforts focused on minimizing the large size and immunogenic potential of Cas9 and other proteins may, in the future, provide complementary tools for the broader mammalian synthetic biology toolkit (*26*, *68*–*70*). Yet, a particularly advantageous feature of ZF-based systems is the ability to quantitatively tune biochemical properties, such as DNA-binding affinity, with simple and general strategies informed by a wealth of biophysical and structural data (*71*, *72*). Indeed, these strategies have been shown to be valuable in designing synthetic regulatory circuits with tunable input/output behaviors (*71*, *73*, *74*), and formed the foundation of our recent efforts to engineer multivalent cooperative transcription factor assemblies in yeast, which we demonstrated to be a compact and flexible mechanism for programming signal processing behaviors in cells (*73*).

Evaluating the true immunogenic potential of any synthetic system will ultimately require empirical measurements in patients. However, a preliminary informatic analysis of our ZFs using an established immunogenicity prediction tool confirms that ZF peptides show comparatively lower immunogenicity scores than those of TetR, Gal4, and *sp* dCas9 (**Methods, Fig. S10**), providing additional evidence that ZFs are overall less likely to elicit immune responses. Indeed, the protein backbone sequences of our engineered ZFs are based entirely on human Egr1; variability is only introduced in residues of the recognition helix of each ZF domain and by altering two residues in the canonical ZF linker to create ‘disrupted’ linkers that connect each 2F unit. Furthermore, our synZiFTRs utilize human effector domains such as p65 to drive strong gene expression outputs in both common laboratory and clinically-relevant human cell types. Recent developments in high-throughput methods to discover and characterize transcriptional effectors should provide additional human-derived effector candidates for artificial transcriptional regulation using our synZiFTR platform (*75*). We recognize that clinical tradeoffs were introduced in our inducible synZiFTR switch designs, as the decision to prioritize clinically-favorable, non-native small molecules required the use of plant-derived ABI-PYL and viral-derived NS3 domains. Future efforts to ‘de-immunize’ these domains will be useful for ultimate clinical translation. Concurrently, efforts to identify additional human ligand binding domains with orthogonal, biocompatible inducers will also be valuable.

We expect that our synZiFTR toolkit will translate widely to other clinically-relevant cell types and contexts, enabling new forms of precise gene expression control. In this study, we focused on establishing small molecule-mediated, remote control of therapeutic activity. Such orthogonal, regulatable gene expression control is urgently needed to improve the safety and efficacy of gene and cell therapies, allowing more potent therapies to be safely administered and controlled post-delivery. Indeed, uncontrolled overexpression of therapeutic genes can lead to toxicity or diminished functionality. For example, poorly controlled IL-12 expression in CAR T cells led to life-threatening side-effects in a recent clinical trial (*76*), and constitutive CAR expression is well-known to cause T cell exhaustion that can limit therapeutic efficacy (*77*). Our work in human cells using xenograft mouse models will pave the way for more in-depth evaluation of toxicity management in immunocompetent mouse models with murine immune cells, which have proven to be more challenging to genetically engineer (*78*). Moreover, gene and cell therapies that perform multiple, concerted therapeutic actions will be critical for addressing complex diseases, including solid cancers, autoimmunity, and heart disease (*62*, *63*). To our knowledge, our study included the first demonstration of a two-channel synthetic switch, based on clinically-approved small molecules, for independent control of therapeutically-relevant genes in primary human immune cells. These results may provide new possibilities to design coengineered cell therapies, which couple the control of factors that direct target specificity with immunomodulatory or immunotherapeutic products to combat different facets of pathology or promote tissue regeneration.

As our synZiFTRs are modular and have mutually-orthogonal specificities, they present a convenient platform for building new gene switches and composing other types of synthetic circuits. On the former, incorporation of *de novo*-designed bioactive protein domains should provide additional modes for regulating the activity of synZiFTRs (*79*–*81*). On the latter, synZiFTR-based circuits that integrate cell-autonomous decision-making will make it possible to activate multi-gene therapeutic programs in response to endogenous and disease-related signaling. To this end, the emergence of flexible synthetic receptors, such as the synthetic Notch receptor, have enabled researchers to endow mammalian cells with novel sense-and-response capabilities to detect a broader set of disease or tissue-related cues and translate them into custom transcriptional outputs (*8*, *9*). In a separate study, we report that synZiFTRs can robustly function within a new class of synthetic transcriptional receptors that undergo regulated intramembrane proteolysis (*82*). This advance allows for activation of custom synZiFTR programs in response to a broad spectrum of ligands via a highly compact, fully humanized receptor system with dramatically expanded input/output capabilities. An exciting future prospect is engineering multi-input synZiFTR circuits that can flexibly integrate information from both exogenously-administered inputs (e.g. small molecules) and cell-autonomous signals (e.g. antigen sensing).

Our synZiFTR toolkit provides a powerful and clinically-promising platform with which to engineer custom transcriptional programs that endow mammalian cells with new capabilities. While much development remains and many other clinical considerations to address, we hope these tools will begin to transform the rapid advances we are witnessing in mammalian synthetic biology into new solutions for safer, effective and powerful next-generation therapies.

## Acknowledgments

We thank Maggie Bobbin for technical assistance with design of zinc finger arrays. We thank C. Bashor, J. Ngo, and members of the Wong and Khalil laboratories for helpful discussions.

## Author contributions

D.V.I. and A.S.K. conceived the study. D.V.I., J.S., and J.K.J. designed ZF proteins and target sequences. D.V.I performed bioinformatic and sequencing analyses. D.V.I. and H.S.L. designed and generated genetic constructs, performed experiments, analyzed the data, and generated figures. K.G. developed 3D tumor spheroid models, and performed spheroid experiments with H.S.L. K.T.R., W.W.W., and A.S.K. oversaw the study and analyzed the data. All authors commented on and approved the manuscript.

## Data and materials availability

All DNA constructs and cell lines are available from A.S.K. All sequencing data is being deposited in the Sequence Read Archive (SRA) and will be accessible with a BioProject accession. Other datasets generated and analyzed during the current study are available upon request from the corresponding author. Computer codes are being made available at github.com/khalillab.

## Supplementary Materials

Materials and Methods

Figures S1-S10

Tables S1-S3

References

## References and Notes

1. M. A. Fischbach, J. A. Bluestone, W. A. Lim, Cell-based therapeutics: the next pillar of medicine. Sci Transl Med 5, 179ps177 (2013).

2. T. Kitada, B. DiAndreth, B. Teague, R. Weiss, Programming gene and engineered-cell therapies with synthetic biology. Science 359, eaad1067 (2018).

3. W. A. Lim, C. H. June, The Principles of Engineering Immune Cells to Treat Cancer. Cell 168, 724–740 (2017).

4. K. A. Hay, Cytokine release syndrome and neurotoxicity after CD19 chimeric antigen receptor-modified (CAR-) T cell therapy. Br J Haematol 183, 364–374 (2018).

5. R. A. Morgan et al., Case report of a serious adverse event following the administration of T cells transduced with a chimeric antigen receptor recognizing ERBB2. Mol Ther 18, 843–851 (2010).

6. G. Fuca, L. Reppel, E. Landoni, B. Savoldo, G. Dotti, Enhancing Chimeric Antigen Receptor T-Cell Efficacy in Solid Tumors. Clin Cancer Res 26, 2444–2451 (2020).

7. X. J. Gao, L. S. Chong, M. S. Kim, M. B. Elowitz, Programmable protein circuits in living cells. Science 361, 1252–1258 (2018).

8. K. T. Roybal et al., Precision Tumor Recognition by T Cells With Combinatorial Antigen-Sensing Circuits. Cell 164, 770–779 (2016).

9. K. T. Roybal et al., Engineering T Cells with Customized Therapeutic Response Programs Using Synthetic Notch Receptors. Cell 167, 419–432 e416 (2016).

10. L. Schukur, B. Geering, G. Charpin-El Hamri, M. Fussenegger, Implantable synthetic cytokine converter cells with AND-gate logic treat experimental psoriasis. Sci Transl Med 7, 318ra201 (2015).

11. M. Xie et al., beta-cell-mimetic designer cells provide closed-loop glycemic control. Science 354, 1296–1301 (2016).

12. F. Sedlmayer, D. Aubel, M. Fussenegger, Synthetic gene circuits for the detection, elimination and prevention of disease. Nat Biomed Eng 2, 399–415 (2018).

13. M. Ptashne, The chemistry of regulation of genes and other things. J Biol Chem 289, 5417–5435 (2014).

14. S. Braselmann, P. Graninger, M. Busslinger, A selective transcriptional induction system for mammalian cells based on Gal4-estrogen receptor fusion proteins. Proc Natl Acad Sci U S A 90, 1657–1661 (1993).

15. M. Gossen, H. Bujard, Tight control of gene expression in mammalian cells by tetracycline-responsive promoters. Proc Natl Acad Sci U S A 89, 5547–5551 (1992).

16. D. Favre et al., Lack of an immune response against the tetracycline-dependent transactivator correlates with long-term doxycycline-regulated transgene expression in nonhuman primates after intramuscular injection of recombinant adeno-associated virus. Journal of virology 76, 11605–11611 (2002).

17. A. M. Lena, P. Giannetti, E. Sporeno, G. Ciliberto, R. Savino, Immune responses against tetracycline-dependent transactivators affect long-term expression of mouse erythropoietin delivered by a helper-dependent adenoviral vector. J Gene Med 7, 1086–1096 (2005).

18. A. Chavez et al., Highly efficient Cas9-mediated transcriptional programming. Nat Methods 12, 326–328 (2015).

19. L. A. Gilbert et al., Genome-Scale CRISPR-Mediated Control of Gene Repression and Activation. Cell 159, 647–661 (2014).

20. L. A. Gilbert et al., CRISPR-mediated modular RNA-guided regulation of transcription in eukaryotes. Cell 154, 442–451 (2013).

21. S. Kiani et al., CRISPR transcriptional repression devices and layered circuits in mammalian cells. Nat Methods 11, 723–726 (2014).

22. L. Nissim, S. D. Perli, A. Fridkin, P. Perez-Pinera, T. K. Lu, Multiplexed and programmable regulation of gene networks with an integrated RNA and CRISPR/Cas toolkit in human cells. Mol Cell 54, 698–710 (2014).

23. P. Perez-Pinera et al., RNA-guided gene activation by CRISPR-Cas9-based transcription factors. Nat Methods 10, 973–976 (2013).

24. J. G. Zalatan et al., Engineering complex synthetic transcriptional programs with CRISPR RNA scaffolds. Cell 160, 339–350 (2015).

25. C. T. Charlesworth et al., Identification of preexisting adaptive immunity to Cas9 proteins in humans. Nat Med 25, 249–254 (2019).

26. S. R. Ferdosi et al., Multifunctional CRISPR-Cas9 with engineered immunosilenced human T cell epitopes. Nat Commun 10, 1842 (2019).

27. A. Mehta, O. M. Merkel, Immunogenicity of Cas9 Protein. J Pharm Sci 109, 62–67 (2020).

28. D. L. Wagner et al., High prevalence of Streptococcus pyogenes Cas9-reactive T cells within the adult human population. Nat Med 25, 242–248 (2019).

29. P. Bai et al., A fully human transgene switch to regulate therapeutic protein production by cooling sensation. Nat Med 25, 1266–1273 (2019).

30. M. Elrod-Erickson, T. E. Benson, C. O. Pabo, High-resolution structures of variant Zif268-DNA complexes: implications for understanding zinc finger-DNA recognition. Structure 6, 451–464 (1998).

31. N. P. Pavletich, C. O. Pabo, Zinc finger-DNA recognition: crystal structure of a Zif268-DNA complex at 2.1 A. Science 252, 809–817 (1991).

32. S. A. Lambert et al., The Human Transcription Factors. Cell 175, 598–599 (2018).

33. V. M. Rivera et al., Long-term pharmacologically regulated expression of erythropoietin in primates following AAV-mediated gene transfer. Blood 105, 1424–1430 (2005).

34. R. R. Beerli, C. F. Barbas, 3rd, Engineering polydactyl zinc-finger transcription factors. Nat Biotechnol 20, 135–141 (2002).

35. M. L. Maeder, S. Thibodeau-Beganny, J. D. Sander, D. F. Voytas, J. K. Joung, Oligomerized pool engineering (OPEN): an ‘open-source’ protocol for making customized zinc-finger arrays. Nat Protoc 4, 1471–1501 (2009).

36. C. O. Pabo, E. Peisach, R. A. Grant, Design and selection of novel Cys2His2 zinc finger proteins. Annu Rev Biochem 70, 313–340 (2001).

37. J. D. Sander et al., Selection-free zinc-finger-nuclease engineering by context-dependent assembly (CoDA). Nat Methods 8, 67–69 (2011).

38. M. Garriga-Canut et al., Synthetic zinc finger repressors reduce mutant huntingtin expression in the brain of R6/2 mice. Proc Natl Acad Sci U S A 109, E3136–3145 (2012).

39. D. Hockemeyer et al., Efficient targeting of expressed and silent genes in human ESCs and iPSCs using zinc-finger nucleases. Nat Biotechnol 27, 851–857 (2009).

40. D. G. Ousterout et al., Correction of dystrophin expression in cells from Duchenne muscular dystrophy patients through genomic excision of exon 51 by zinc finger nucleases. Mol Ther 23, 523–532 (2015).

41. V. Sebastiano et al., In situ genetic correction of the sickle cell anemia mutation in human induced pluripotent stem cells using engineered zinc finger nucleases. Stem cells 29, 1717–1726 (2011).

42. F. D. Urnov et al., Highly efficient endogenous human gene correction using designed zinc-finger nucleases. Nature 435, 646–651 (2005).

43. J. Zou et al., Gene targeting of a disease-related gene in human induced pluripotent stem and embryonic stem cells. Cell Stem Cell 5, 97–110 (2009).

44. Y. Choo, I. Sanchez-Garcia, A. Klug, In vivo repression by a site-specific DNA-binding protein designed against an oncogenic sequence. Nature 372, 642–645 (1994).

45. L. Zhang et al., Synthetic zinc finger transcription factor action at an endogenous chromosomal site. Activation of the human erythropoietin gene. J Biol Chem 275, 33850–33860 (2000).

46. M. Moore, A. Klug, Y. Choo, Improved DNA binding specificity from polyzinc finger peptides by using strings of two-finger units. Proc Natl Acad Sci U S A 98, 1437–1441 (2001).

47. J. K. Rockstroh et al., Efficacy and safety of grazoprevir (MK-5172) and elbasvir (MK-8742) in patients with hepatitis C virus and HIV co-infection (C-EDGE CO-INFECTION): a non-randomised, open-label trial. Lancet HIV 2, e319–327 (2015).

48. C. L. Jacobs, R. K. Badiee, M. Z. Lin, StaPLs: versatile genetically encoded modules for engineering drug-inducible proteins. Nat Methods 15, 523–526 (2018).

49. E. P. Tague, H. L. Dotson, S. N. Tunney, D. C. Sloas, J. T. Ngo, Chemogenetic control of gene expression and cell signaling with antiviral drugs. Nat Methods 15, 519–522 (2018).

50. R. Feil et al., Ligand-activated site-specific recombination in mice. Proc Natl Acad Sci U S A 93, 10887–10890 (1996).

51. A. K. Indra et al., Temporally-controlled site-specific mutagenesis in the basal layer of the epidermis: comparison of the recombinase activity of the tamoxifen-inducible Cre-ER(T) and Cre-ER(T2) recombinases. Nucleic Acids Res 27, 4324–4327 (1999).

52. F. S. Liang, W. Q. Ho, G. R. Crabtree, Engineering the ABA plant stress pathway for regulation of induced proximity. Sci Signal 4, rs2 (2011).

53. H. K. Chung et al., Tunable and reversible drug control of protein production via a self-excising degron. Nat Chem Biol 11, 713–720 (2015).

54. J. Cohen, IL-12 deaths: explanation and a puzzle. Science 270, 908 (1995).

55. J. P. Leonard et al., Effects of single-dose interleukin-12 exposure on interleukin-12-associated toxicity and interferon-gamma production. Blood 90, 2541–2548 (1997).

56. P. Chevallier et al., Trastuzumab for treatment of refractory/relapsed HER2-positive adult B-ALL: results of a phase 2 GRAALL study. Blood 119, 2474–2477 (2012).

57. P. Chevallier et al., Overexpression of Her2/neu is observed in one third of adult acute lymphoblastic leukemia patients and is associated with chemoresistance in these patients. Haematologica 89, 1399–1401 (2004).

58. S. P. Haen et al., Prognostic relevance of HER2/neu in acute lymphoblastic leukemia and induction of NK cell reactivity against primary ALL blasts by trastuzumab. Oncotarget 7, 13013–13030 (2016).

59. S. K. Joshi et al., ERBB2/HER2 mutations are transforming and therapeutically targetable in leukemia. Leukemia 34, 2798–2804 (2020).

60. J. H. Cho, J. J. Collins, W. W. Wong, Universal Chimeric Antigen Receptors for Multiplexed and Logical Control of T Cell Responses. Cell 173, 1426–1438 e1411 (2018).

61. H. P. Feng et al., Pharmacokinetic Interactions between the Hepatitis C Virus Inhibitors Elbasvir and Grazoprevir and HIV Protease Inhibitors Ritonavir, Atazanavir, Lopinavir, and Darunavir in Healthy Volunteers. Antimicrob Agents Chemother 63, (2019).

62. E. Lanitis, G. Coukos, M. Irving, All systems go: converging synthetic biology and combinatorial treatment for CAR-T cell therapy. Curr Opin Biotechnol 65, 75–87 (2020).

63. S. Mardiana, B. J. Solomon, P. K. Darcy, P. A. Beavis, Supercharging adoptive T cell therapy to overcome solid tumor-induced immunosuppression. Sci Transl Med 11, (2019).

64. M. Hong, J. D. Clubb, Y. Y. Chen, Engineering CAR-T Cells for Next-Generation Cancer Therapy. Cancer Cell 38, 473–488 (2020).

65. M. Liu et al., TGF-beta suppresses type 2 immunity to cancer. Nature 587, 115–120 (2020).

66. F. Mittler et al., High-Content Monitoring of Drug Effects in a 3D Spheroid Model. Front Oncol 7, 293 (2017).

67. S. Herter et al., A novel three-dimensional heterotypic spheroid model for the assessment of the activity of cancer immunotherapy agents. Cancer Immunol Immunother 66, 129–140 (2017).

68. E. Kim et al., In vivo genome editing with a small Cas9 orthologue derived from Campylobacter jejuni. Nat Commun 8, 14500 (2017).

69. D. Ma, S. Peng, W. Huang, Z. Cai, Z. Xie, Rational Design of Mini-Cas9 for Transcriptional Activation. ACS Synth Biol 7, 978–985 (2018).

70. F. A. Ran et al., In vivo genome editing using Staphylococcus aureus Cas9. Nature 520, 186–191 (2015).

71. A. S. Khalil et al., A Synthetic Biology Framework for Programming Eukaryotic Transcription Functions. Cell 150, 647–658 (2012).

72. J. C. Miller et al., Enhancing gene editing specificity by attenuating DNA cleavage kinetics. Nat Biotechnol 37, 945–952 (2019).

73. C. J. Bashor et al., Complex signal processing in synthetic gene circuits using cooperative regulatory assemblies. Science 364, 593–597 (2019).

74. P. S. Donahue et al., The COMET toolkit for composing customizable genetic programs in mammalian cells. Nat Commun 11, 779 (2020).

75. J. Tycko et al., High-Throughput Discovery and Characterization of Human Transcriptional Effectors. Cell 183, 2020–2035 e2016 (2020).

76. L. Zhang et al., Tumor-infiltrating lymphocytes genetically engineered with an inducible gene encoding interleukin-12 for the immunotherapy of metastatic melanoma. Clin Cancer Res 21, 2278–2288 (2015).

77. A. H. Long et al., 4-1BB costimulation ameliorates T cell exhaustion induced by tonic signaling of chimeric antigen receptors. Nat Med 21, 581–590 (2015).

78. E. Lanitis et al., Optimized gene engineering of murine CAR-T cells reveals the beneficial effects of IL-15 coexpression. J Exp Med 218, (2021).

79. R. A. Langan et al., De novo design of bioactive protein switches. Nature 572, 205–210 (2019).

80. A. H. Ng et al., Modular and tunable biological feedback control using a de novo protein switch. Nature 572, 265–269 (2019).

81. Z. Chen et al., Programmable design of orthogonal protein heterodimers. Nature 565, 106–111 (2019).

82. I. Zhu et al., Comprehensive clinically driven design of synthetic receptors for custom gene circuit regulation in therapeutic cells. In preparation.

