## Supplementary Materials for "Clinically-driven design of synthetic gene regulatory programs in human cells"

**This PDF files includes:**

Materials and Methods
Figs. S1 to S10
Tables S1 to S3
References

#### Materials and Methods

##### *Zinc Finger Engineering*

Design and assembly of ZF arrays. Engineered six-finger arrays and their corresponding DNA-binding motifs (DBMs) were designed using a proprietary archive of preselected two-finger (2F) subarrays that recognize >600 unique 6-bp DNA subsites (**Fig. S1C**). Each 2F subarray, composed of two tandemly linked ZFs, was co-selected in the context of a 3F array to recognize 6-bp of DNA. Libraries of 2F subarrays were developed from highly functional 3F arrays experimentally characterized using previously described bacterial-2-hybrid (B2H) approaches (*1-3*). Six-finger arrays were created by modularly connecting three 2F units separated by 'disrupted' linkages, to disfavor cooperative binding of adjacent units to potentially nonspecific sequences (*4-9*) (**Fig S1B**). Where possible, a context-dependent approach considering adjacent 5' and 3' nucleotides was applied to choose 2F subarray combinations that would likely retain their DNA binding performances in higher-order arrays.

Determination of targetable, genome-orthogonal DNA-binding motifs. Concatenating three 6-bp subsites generates 18-bp core recognition sequences that are targetable by six-finger ZF arrays. To nominate target sequences that are "orthogonal" to the human genome, we used the following design criteria. 1) Target sequences were composed predominately of 6-bp subsites that were informatically determined to occur relatively infrequently in the human genome **(Fig S1D)**. Concatenating these infrequent subsites creates longer motifs that are unlikely to be found in the genome by random chance. 2) Candidate 18-bp sequences were chosen to minimize identity from exact or near-matched sequences found in the reference human genome while maintaining a balanced composition **(Fig 1C)**. 3) ZF arrays are known to exhibit preferential affinity to particular 5' and 3' flanking base pairs (*10, 11*). Thus, their recognition specificities were effectively increased to 20-bp sequences, which we termed 'DNA-binding motifs' (DBMs).

##### *Bioinformatic Analysis of Genomic Sequence Occurrence*

Computational sequence analysis of DBMs was performed using Biostrings, an open-source Bioconductor R software package containing efficient string alignment algorithms (https://bioconductor.org/). The UCSC hg19 reference human genome sequence was obtained from a Bioconductor Annotation Package. Custom scripts to count exact or mismatched strings, as well as to return information on corresponding genomic locations, were developed based on open-source Bioconductor vignettes for Genome Searching. As the hg19 genome object only contained the forward strand sequence, analysis of each input sequence involved a summation of outputs from both the forward and reverse complement of the sequence. Sequences of response elements for common artificial regulators (Gal4 UAS, TetO, ZFHD1) were obtained from the literature (*12*). All of the DBM sequences used in this analysis can be found in **Table S1**.

##### *Bioinformatic Analysis of Predicted Immunogenicity*

Computational sequence analysis of DNA-binding domain sequences was performed using the T Cell Class I pMHC Immunogenicity Tool, an open-source tool from the Immune Epitope Database used to predict the relative chance a peptide/MHC complex will elicit an immune response (*13*). Sequences were processed using IEDB-Recommended default settings. Sequences of DNA-binding domains for commonly-used regulators were obtained from the literature (Gal4, TetR, *Streptococcus pyogenes* Sp-dCas9) (*12*). All of the protein sequences used in this analysis can be found in **Table S2**.

##### *Plasmid Design and Construction*

All plasmids used in this study are listed in **Table S3**. Plasmids were constructed using established molecular biology methods and Gibson isothermal assembly. Engineered cassettes were subcloned into vectors containing Ampicillin resistance as a bacterial selection marker. All plasmids were sequence-verified and archived in competent *E. coli* TG1 (GoldBio CC-205-A). Transient transfection plasmids were constructed by subcloning cassettes into the pCAGEN mammalian expression vector backbone (Addgene #11160) digested with SalI/HindIII. Donor plasmids for CRISPR-Cas9 mediated AAVS1 locus integration were constructed by subcloning cassettes into the pMP472 vector backbone digested with NgoMIV/AscI (*14*); these vectors were additionally modified to contain Puromycin resistance and BFP, separated by a 2A sequence, as mammalian selection markers. Donor plasmids for lentiviral integration were constructed by subcloning cassettes into the pHR'SIN vector backbone digested with EcoRI/NotI (*15*).

Response promoter cassettes contained arrays of either 4 or 8 copies of DBMs, each separated by neutral 18-bp DNA spacers designed using the open-source program R2oDNA Designer (*16*). Additional 39-bp DNA spacers, designed using R2oDNA Designer, were used to flank the 5' and 3' ends of each DBM array and function as conserved insulation sequences. Each DBM array was cloned upstream of a minimal promoter (minCMV, minTK, ybTATA) or a full-length promoter (pSFFV). SynZiFTR expression cassettes contained a constitutive promoter (pUb, pSFFV) driving expression of either synZiFTR or 'mock' negative controls. Minimal synZiFTRs contained a 5' 3X FLAG tag followed by a SV40 nuclear localization signal. All inducible synZiFTRs contained a SV40 NLS, with the exception of ERT2 fusions. Bicistronic cassettes used for RNA-seq and single lentiviral transductions contained (5'-3') response promoter cassettes in the reverse orientation, followed by synZiFTR expression cassettes in the forward orientation.

##### *Mammalian Cell Culture*

All cells were maintained in 10 cm treated dishes (adherent) or 75 mL flasks (suspension) at 37C and 5% CO2. Cells were passaged every 3-4 days when they reached 70-80% confluency or ~2 million cells/mL. Cell lines were not used for experiments beyond 40 passages.

HEK293FT cell lines (Thermo Fisher Scientific R700-07) were cultured in Dulbecco's Modified Eagle's Medium (Corning 10-013-CV) supplemented with 10% Fetal Bovine Serum

(Takara Bio 631367), 1% GlutaMAX (Thermo Scientific 35050061), 1% Non-Essential Amino Acids (Thermo Scientific 11140050), and 1% Penicillin-Streptomycin (Thermo Scientific 15140122). Jurkat cell lines (ATCC TIP-152) were cultured in RPMI 1640 Medium (Corning 10-040-CV) supplemented with 10% FBS, 1% GlutaMAX, and 1% Pen-Strep.

NALM6 target cell lines were cultured in RPMI 1640 Medium supplemented with 5% FBS, 1% L-glutamine (Gibco A2916801), and 1% Pen-Strep. MCF10A target cell lines were cultured in DMEM/F-12 (Gibco 11320033) supplemented with 5% Horse Serum (Gibco 16050130), 1% Pen-Strep, 20ng/mL EGF (Peprotech), 0.5 mg/mL hydrocortisone (Sigma H-0888), 100ng/mL Cholera Toxin (Sigma C-8052), and 10ug/mL Insulin (Sigma I-1882).

Primary peripheral blood mononuclear cells (PBMC) were obtained from whole peripheral blood from healthy donors at the Blood Donor Center at Boston Children's Hospital through a protocol approved by the Boston University Institutional Review Board (IRB). PBMCs were isolated using Lymphoprep (STEMCELL Technologies NC0418243). Cells were cultured in X-Vivo 15 Medium (Lonza 04-418Q) supplemented with 5% human AB serum (Valley Biomedical, Inc. HP1022), 10mM N-acetyl L-Cysteine (Sigma A9165), 55 uM 2-Mercaptoethanol (Gibco), and 100-200 U / mL IL-2 (NCI BRB Preclinical Repository).

***Transient Transfection***

Plasmids were transiently transfected into HEK293FT cell lines using polyethylenimine (PEI, 7.5 mM linear PEI stock, nitrogen/phosophorus ratio of 20, Polysciences). 40,000 HEK293FT cells were seeded into 96-well treated plates for 24 hours and subsequently transfected with 300 ng total DNA in a NaCl-PEI solution. All constructs were diluted to 100 ng/uL in deionized water prior to being added to the transfection mix. The total amount of transfected DNA contained an equal ratio of all desired constructs, including a constitutive iRFP-expressing plasmid as a transfection marker and salmon sperm DNA as a filler. Upon transfection, cells were incubated for 48-72 hours prior to downstream flow cytometry measurements. All transfections were performed in biological triplicate.

***Stable Cell Line Generation Using CRISPR/Cas9-Mediated Integration***

Cassettes cloned into CRISPR/Cas9 donor plasmids were integrated into HEK293FT cell lines using CRISPR/Cas9 mediated integration into the AAVS1 (PPP1R2C) safe-harbor locus (*14*). 500,000 HEK293FT cells were seeded into 6-well treated plates for 24 hours and subsequently transfected with 2 ug total DNA in a NaCl-PEI solution. All constructs were diluted to 100 ng/uL in deionized water prior to being added to the transfection mix. 1 ug of the donor plasmid was co-transfected with 500 ng of the VP12 humanSpCas9-Hf1 plasmid (Addgene #72247) and 500 ng of the gRNA_AAVS1-T2 plasmid (Addgene #11160). Upon transfection, cells were incubated for 72 hours prior to downstream selection with media containing 2 ug/mL Puromycin. Cells were selected for at least 3 passages prior to transfection or RNA sequencing experiments.

##### ***Stable Cell Line Generation Using Lentiviral Integration***

Cassettes cloned into lentiviral donor plasmids were integrated into Jurkat cell lines using lentiviral infection. Lentivirus was harvested from transfected HEK293FT cells; 500,000 HEK293FT cells were seeded into 6-well treated plates for 24 hours and subsequently transfected with 2 ug total DNA in a NaCl-PEI solution. All constructs were diluted to 100 ng/uL in deionized water prior to being added to the transfection mix. 1 ug of the donor plasmid was co-transfected with 700 ng of the pCMVR8.74 plasmid (Addgene #22036), 100 ng of the pAdVantage plasmid (Promega), and 200 ng of the pMD2.G VSVG plasmid (Addgene #12259). Upon transfection, cells were incubated for 72 hours prior to harvesting media containing lentiviral supernatant. Lentiviral media was centrifuged for 5 minutes at 300 g. 500,000 Jurkat cells were seeded into 12-well treated plates in 1 mL media. 1 mL of each lentiviral media was added to the cells (approximate MOI = 30). Infected cells were incubated for 24-48 hours prior to removal of lentivirus. Cells were collected and centrifuged for 5 minutes at 300 g. Lentiviral media was removed and fresh media was added. Transduced cells were subsequently used for downstream induction measurements.

Cassettes cloned into lentiviral donor plasmids were also integrated into primary PBMCs using lentiviral infection. Following 24 hours HEK293FT co-transfection, media was replaced with UltraCULTURE (Lonza 12-725F) supplemented with Pen-Strep, L-glutamine, 0.5M Sodium Butyrate and 1mM Sodium Pyruvate. Cells were incubated for 48 hours prior to harvesting media containing lentiviral supernatant. Lentiviral media was centrifuged for 5 minutes at 300 g to remove cell debris, and then concentrated at a 3:1 ratio, either using Lenti-X Concentrator (Takara Bio 631232) or Lentivirus concentration solution (40% PEG8000 and 1.5M NaCl). Cells were incubated at 4C overnight, prior to centrifugation at 4C at 1500 g for 45 minutes (Lenti-X) or 1600 g for 60 minutes (concentration solution). Lentiviral spinfection was performed to transduce primary PBMCs. Cells were thawed 48 hours prior to spinfection, and activated using Human T-activator CD3/CD28 Dynabeads (Thermo Fisher 11131D) with 200U/ml IL-2 media 24 hours prior to spinfection. Concentrated lentivirus was plated on non-TC treated 6-well plates coated with retronectin (Takara Bio T100B) and spun for 90 minutes at 1200xg. Lentivirus was then aspirated and activated T cells were added to the virus-coated wells. Transduced cells were incubated at 37C.

##### ***Flow Cytometry***

All flow cytometry measurements were performed on an Attune NxT Flow Cytometer (Thermo Fisher Scientific). Cell samples were suspended in 200 uL fresh culture media and measured on the flow cytometer in biological triplicate. Live cells were gated by forward scatter (FSC) and side scatter (SSC). Fluorescence data was collected for GFP (excitation laser: 488nm, emission filter: 530/30nm), iRFP-720 (excitation laser: 640nm, emission filter: 720/30nm), BFP (excitation laser: 405nm, emission filter: 440/50nm), and mCherry (excitation laser: 561nm, emission filter: 620/15nm). A minimum of 10,000 live cells were collected for each sample. Flow cytometry data was analyzed using FlowJo (Treestar Software). Live cells were gated by forward scatter and side scatter. Transfected cells were gated for the presence of the iRFP transfection marker. CRISPR-integrated cells were gated for the presence of the BFP integration marker. Geometric means of fluorescence distributions were calculated by FlowJo. Data was further analyzed using the Prism 8 software (GraphPad).

##### *Fluorescence Activated Cell Sorting (FACS)*

FACS was performed for HEK293FT cell lines stably integrated with bicistronic cassettes on a SH800 Cell Sorter (Sony Corporation). Cell samples were suspended in 2 mL 1X Phosphate Buffered Saline containing 1% Fetal Bovine Serum and passed through a 0.45 um filter to break clumps. Live cells were gated by forward scatter (FSC) and side scatter (SSC). Fluorescence data was collected for GFP (excitation laser: 488nm) and mCherry (excitation laser: 561nm). Individual clones of ‘reference’ and ‘test’ lines highly expressing GFP or mCherry were respectively sorted into 96-well treated plates containing 200 uL fresh culture media. Clones were incubated for approximately 14 days until several clones reached sufficient population density. Populations were measured on the Attune NxT Flow Cytometer using the aforementioned flow cytometry protocol to verify expression of GFP and mCherry, and a single population for each condition was maintained for RNA sequencing.

FACS was also performed for primary PBMCs stably integrated with cassettes. Cell samples were suspended in 3 mL fresh PBMC culture media. Live cells were gated by forward scatter (FSC) and side scatter (SSC). Fluorescence data was collected for either mCherry (excitation laser: 561nm) or anti-tEGFR(AF647) (excitation laser: 638nm). Gated populations of cells highly expressing mCherry (CAR+) or tEGFR (payload+) were respectively sorted into tubes at a concentration of 1 million cells / mL. Sorted cells were activated using Human T-activator CD3/CD28 Dynabeads (Thermo Fisher Scientific) at 1:2 cell-to-bead ratio and cultured in fresh media containing 100U/mL IL2 for approximately 7 days.

##### *RNA Sequencing*

All RNA sequencing preparation steps and measurements were performed in duplicate beginning with total RNA collection. Total RNA was purified from 1-2 million of each sorted HEK293FT population using the RNeasy Plus Mini Kit (Qiagen) and eluted to a final concentration of 200-500 ng/uL. Samples were submitted to the Tufts University Core Facility (TUCF Genomics) for subsequent preparation of sequencing libraries. Sample purity was measured by TUCF Genomics using an Agilent Bioanalyzer; cDNA synthesis and libraries were prepared using the TruSeq Stranded mRNA Library Prep Kit (Illumina). These libraries were sequenced as 50 bp Single End reads using a HiSeq 2500 instrument (Illumina).

Sequences were analyzed against the hg19 reference human genome (*17*). Indices for the reference genome and custom sequences for non-native transcripts (i.e. synZiFTRs, ‘mock’ TFs, and mCherry) were built using the Bowtie2 software (*18*). Transcript reads were aligned to genomic builds using the TopHat software (*19*). Features were annotated with the hg19 GRCh37.p13 annotation file (Gencode) and custom annotation files for non-native transcripts using the featureCounts software. Transcript length- and sequencing depth- normalized counts of individual transcripts were calculated using the following equation for 'Transcripts per Kilobase Million' (TPM):

$$TPM\ normalized\ count\ of\ gene\ X = 10^6 * \frac{\frac{avg.count\ for\ gene\ X}{length\ of\ gene\ X}}{sum(\frac{avg.count\ for\ all\ genes}{length\ of\ all\ genes})}$$

Correlation plots were generated using MATLAB (Mathworks). Differential analysis using the outputs from featureCounts was performed in R using the DESeq2 package (*20*). DESeq2 employed the Benjamini-Hochberg multiple hypothesis correction with a False Discovery Rate cutoff of 5%.

##### *Small Molecule Induction*

Stock solutions of Abscisic Acid (Sigma Aldrich, 50 mM in ethanol), 4-hydroxytamoxifen (Sigma Aldrich, 1 mM in ethanol), and grazoprevir (MedChemExpress, 1 mM in DMSO) were stored at -80C. 50,000 Jurkat cells or primary PBMCs were seeded into 96-well untreated round-bottom plates in 100 uL fresh media. 100 uL of media containing 2X concentrated amounts of the small molecule inducer was added to each well on day 0 of induction. For longer time course experiments, cells were passaged in refreshed induction media every 2-4 days. All inductions were performed in biological triplicate.

##### *Enzyme-Linked Immunosorbent Assay (ELISA)*

ELISA was performed with supernatant from cultured cells using OptEIA kits to measure human IFN-γ (BD Biosciences 555142), human IL-4 (BD Biosciences 555194), or human IL-12(p70) (BD Biosciences 555183). ELISA measurements utilized 0.05% Tween-20 in PBS wash buffer (Thermo Scientific 28352), supplemental reagents from OptEIA Reagent Set B (BD Biosciences 550534), and 96-well flat-bottom MaxiSorp plates (Thermo Fisher Scientific 442404). Absorbance was measured at 450 nm using a SpectraMax microplate reader (Molecular Devices). All ELISA measurements were performed in biological triplicate.

##### *In Vitro Co-Culture Experiments*

Primary PBMCs were co-cultured with Her2+ or CD19+ NALM6 target cells at a 1:1 effector to target (E:T) ratio overnight at 37C. Following 16 hours of incubation, supernatant was saved for ELISA and cells were analyzed using flow cytometry to count live NALM6 cells by gating for BFP+ cells. Killing efficiency was calculated as the percentage of cells killed compared to control wells containing NALM6 cells without PBMCs.

##### *Mouse Xenograft Model Experiments*

Female NOD.Cg-Prkdcscid Il2rgtm1Wjl/SzJ (NSG) mice, 6-8 weeks of age, were purchased from Jackson Laboratories (#005557) and maintained in the BUMC Animal Science Center (ASC). All protocols were approved by the Institutional Animal Care and Use Committee at BUMC. To generate the intravenous blood tumor xenograft models, NSG mice were initially injected with 0.5 X $10^6$ luciferized Her2+ NALM6 cells intravenously. After 3 days, 15 X $10^6$ PBMC cells were infused intravenously. For GZV treatment, Grazoprevir potassium salts (MedChemExpress, HY-15298A) were dissolved in 2.5% DMSO, 30% PEG400 and 67.5% PBS and intraperitoneally injected every day at a dose of 25 mg/kg. For the GZV + LPV/RTV treatment, Lopinavir (MedChemExpress, ABT-378) and Ritonavir (MedChemExpress, ABT-538) were formulated in a 4:1 weight ratio using a previously described strategy and co-administered with 25 mg/kg GZV at a dose of 10 mg/kg (*21*). Tumor burden was measured by IVIS Spectrum (Xenogen) and was quantified as total flux (photons per sec) in the region of interest. Images were acquired within 10 minutes following intraperitoneal injection of 150 mg/kg of D-luciferin (PerkinElmer #122799).

***3D Tumor Spheroid Model Experiments***

Non-adherent spheroid preparation wells were prepared by adding 70 uL of pre-warmed 1% agarose in PBS to each well of a 96-well plate and allowing the wells to cool down for 1 hour. To generate 3D spheroids, 10,000 cells were seeded into preparation wells for 5 days and subsequently transferred into an opaque glass-bottomed dish by pipetting with a cut 200 uL tip. Primary PBMCs were co-cultured with MCF10A target spheroids at a 1:2 effector to target (E:T) ratio in 200 uL media containing 1 uM concentration of small molecule inducers. Following 48 hours of incubation at 37C on a rocking plate, spheroid morphology and PBMC fluorescence images were captured using a Nikon TE200 brightfield microscope equipped with a 10x objective and SPOT insight CMOS camera. 100 uL of supernatant from each well was saved for IL-12 ELISA, which was performed as described above.

***Quantification and Statistical Analysis***

Data between two groups was compared using an unpaired two-tailed t-test as indicated; data between three or more groups was compared using one-way ANOVA with Dunnett's Multiple Comparisons post-hoc test or two-way ANOVA with Tukey's Multiple Comparisons post-hoc test as indicated. All statistical analyses were performed with Prism 9 (GraphPad) and p values are reported (not significant (ns): $p>0.05$, *: $p<0.05$, **: $p<0.01$, ***: $p< 0.001$, ****: $p<0.0001$). All error bars are represented either SEM or SD.

### Supplementary Figures

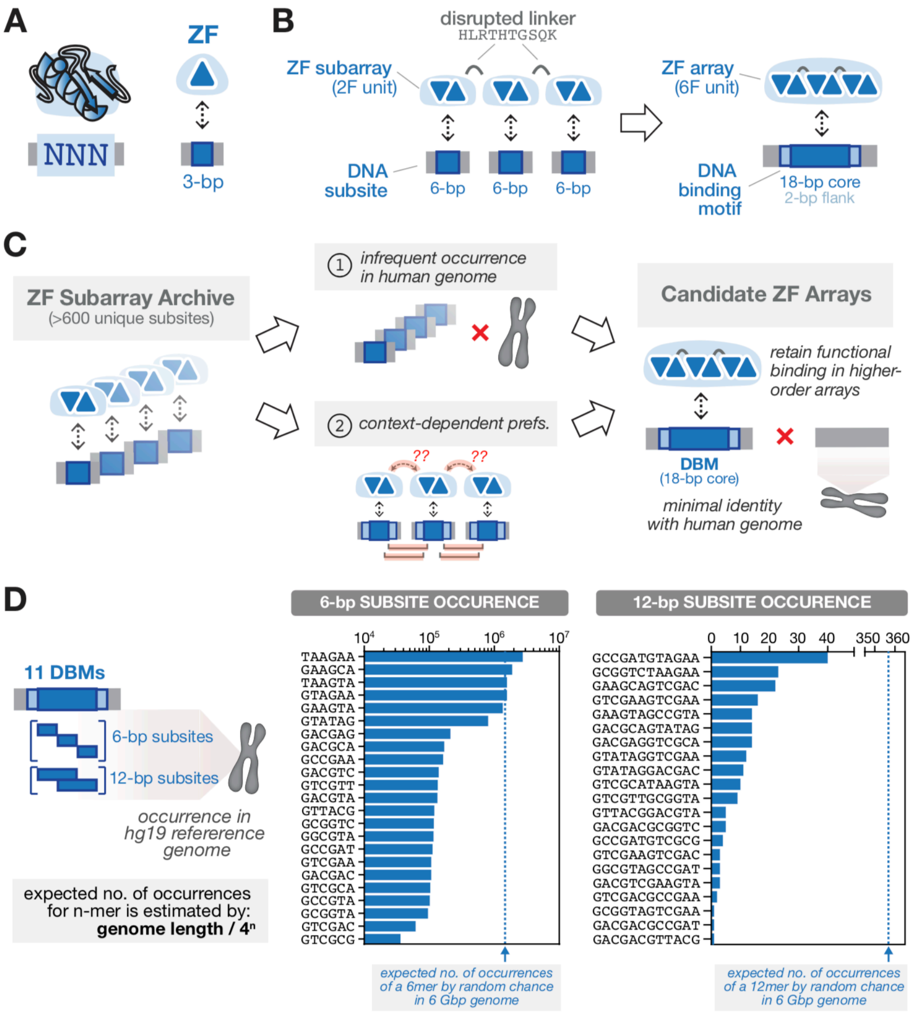


**Fig. S1. Designing ZF arrays that target human genome-orthogonal sequences**
(**A**) Cys2-His2 zinc fingers (ZFs) are small (~30 amino acid) domains that recognize ~3-bp DNA sequences. (**B**) Design and assembly of ZF arrays. By linking two-finger units (each recognizing 6-bp subsites) using flexible 'disrupted' linkers, it is possible to construct functional six-finger arrays capable of specifically recognizing 20-bp DBM sequences (18-bp core + 2-bp flanking). (**C**) Workflow for engineering ZF arrays with genome-orthogonal specificities. From a large archive of pre-selected 2F subarrays, we chose 6-bp subsites that occur infrequently in the human genome, and considered context-dependent preferences for adjacent 2F units. These building blocks lead to a set of DBMs that minimize identity with the human genome, and assembly of 6F arrays that retain functional binding in the concatenated form. (**D**) Prevalence of DBM subsites in the human genome. Occurrences of 6-bp (left) and 12-bp (right) subsites used in the design of full DBMs. The expected numbers of occurrences by random chance in the human genome are indicated by dashed lines.

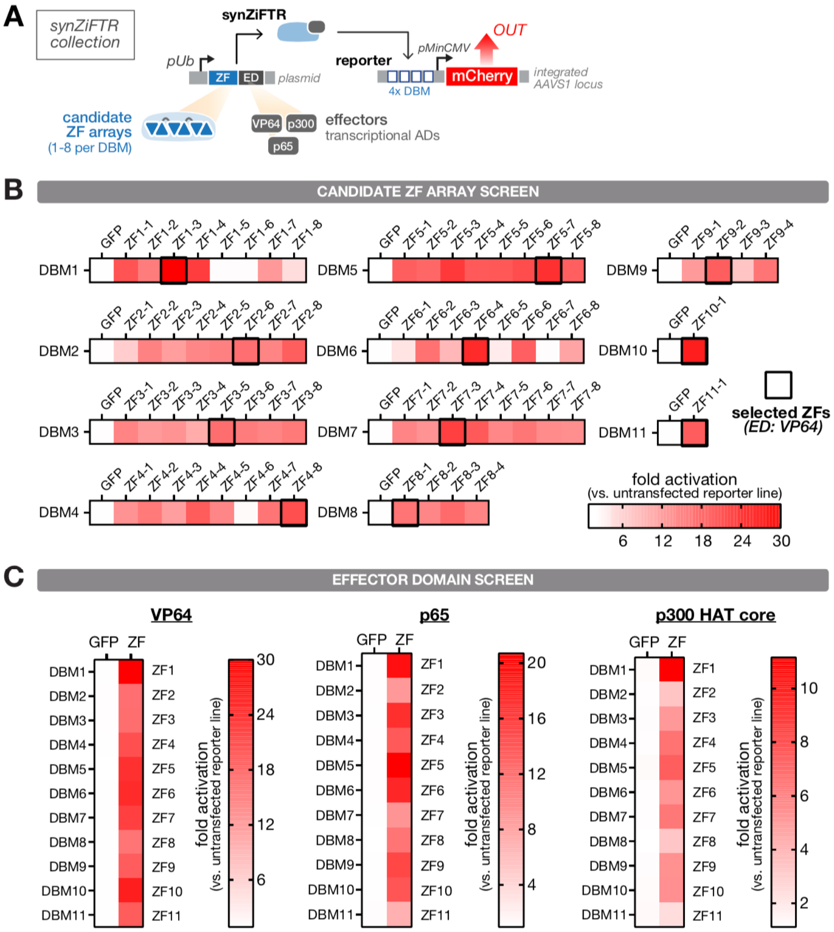


**Fig. S2. Screening synZiFTR domain variants**

(**A**) Schematic of genetic circuit used to screen synZiFTR domain variants: (1) candidate ZF arrays for each DBM and (2) transcriptional activation domains (ADs). Response element vectors for each DBM were stably-integrated into the *AAVS1* locus of HEK293FT cells to generate reporter lines. SynZiFTR (or control) expression vectors were transfected into corresponding reporter lines, and mCherry was measured by flow cytometry after 2 days. (**B**) Screening candidate ZF arrays for each DBM. For each DBM, the ZF that yielded the highest reporter fold activation level was selected (black boxes). Fold activation levels represent mean values for three measurements. (**C**) Screening synZiFTR activity for different transcriptional effector domains: VP64, p64, p300. Fold activation levels represent mean values for three biological replicates.

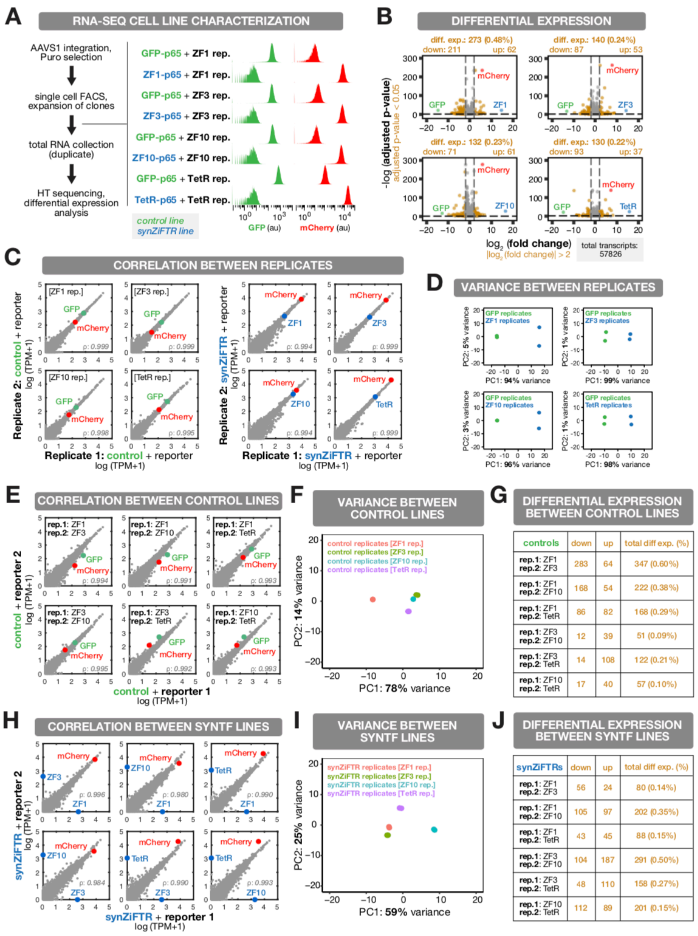


**Fig. S3. Genome-wide transcriptional specificity of synZiFTRs**

(**A**) Overview of RNA-sequencing workflow and cell line characterization. Response elements and corresponding synZiFTRs (or controls) were stably-integrated into HEK293FT cells. Histograms represent fluorescence measurements of GFP (control TF) and mCherry (reporter) from sorted cell lines, prior to total RNA collection in duplicate. (**B**) Differential expression analysis between synZiFTR and control lines. Points represent individual transcripts. Dashed lines indicate the thresholds used to determine differential expression, i.e. significant fold change (|log2 FC| >2) and False Discovery Rate (padj < 0.05). Transcripts identified as differentially expressed are indicated in gold. The total numbers of differentially expressed transcripts (upregulated and downregulated), as well as the corresponding percentage of total transcripts, are indicated above each plot. (**C-D**) Pairwise comparisons of transcript expression between technical replicate samples. Plots represent correlations of normalized transcript abundances between technical replicates (C). Points represent individual transcript levels normalized to TPM, transcripts per kilobase million. Pearson correlation coefficient was calculated for native (grey) transcripts. Overall variance between technical replicates for synZiFTR line samples and corresponding control line samples was measured using principal component analysis (D). (**E-G**) Pairwise comparisons between different 'control' lines. Plots represent correlations of normalized transcript abundances between GFP-p65 expressing lines, indicated by their co-integrated reporters (E). Points represent individual transcript levels normalized to TPM. Pearson correlation coefficient was calculated for native (grey) transcripts. Overall variance between each control replicate sample was measured using principal component analysis (F). Differential expression analysis between pairwise comparisons are summarized in the table (G). Thresholds to determine significant fold change (|log2 FC| >2) and False Discovery Rate (padj < 0.05) were applied to evaluate differential expression. (**H-J**) Pairwise comparisons between different 'synZiFTR' lines. Plots represent correlations of normalized transcript abundances between synZiFTR expressing lines, indicated by the corresponding synZiFTR transcript (H). Points represent individual transcript levels normalized to TPM. Pearson correlation coefficient was calculated for native (grey) transcripts. Overall variance between each synZiFTR replicate sample was measured using principal component analysis (I). Differential expression analysis between pairwise comparisons are summarized in the table (J). Thresholds to determine significant fold change (|log2 FC| >2) and False Discovery Rate (padj < 0.05) were applied to evaluate differential expression.

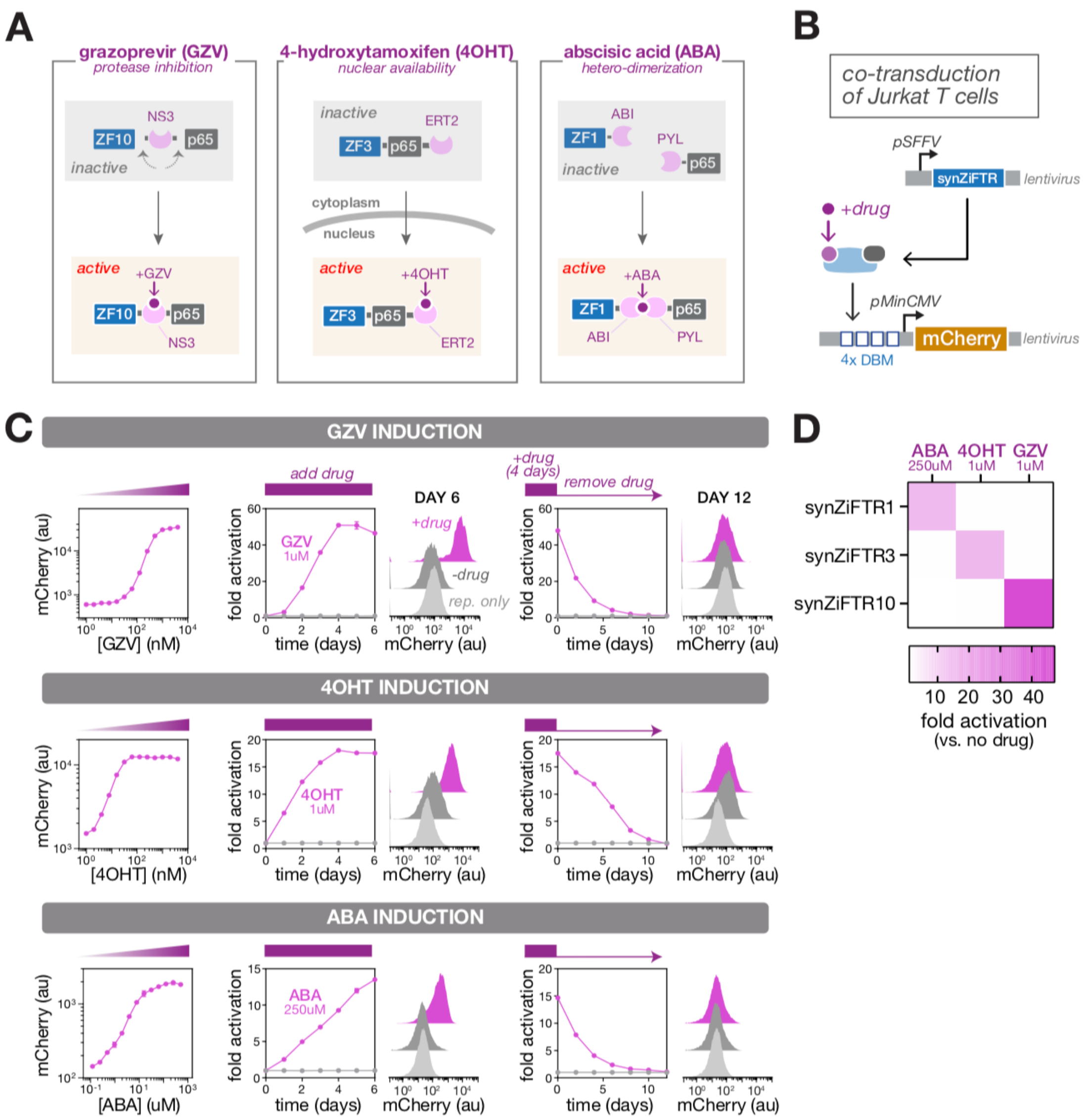


**Fig. S4. Characterization of small molecule-regulated synZiFTR switches**

(**A**) Design of three orthogonal synZiFTR switches, controlled by grazoprevir (GZV), 4-hydroxytamoxifen (4OHT), and abscisic acid (ABA). NS3, hepatitis C virus NS3 protease domain; ERT2, human estrogen receptor T2 mutant domain; ABI, ABA-insensitive 1 domain (aa 126-423); PYL, PYR1-like 1 domain (aa 33-209). (**B**) Schematic of genetic circuit used to characterize small molecule-regulated synZiFTR switches in Jurkat T cells. Jurkat T cells were co-transduced with reporter and synZiFTR expression lentiviral vectors in an equal ratio. (**C**) Dose- and time-responses for the three synZiFTR switches. Dose-responses were generated by measuring mCherry by flow cytometry 4 days following induction with different concentrations of each small molecule inducer (left). Time-responses were generated by inducing cells with saturating concentrations of each inducer, and measuring mCherry at defined time points (center). De-activation dynamics following removal of inducer (right). Points represent mean values for three measurements ± SD. Histograms show absolute levels for one representative measurement. (**D**) Small molecule inducers specifically activate respective synZiFTR switches. Fold activation levels represent mean values for three biological replicates.

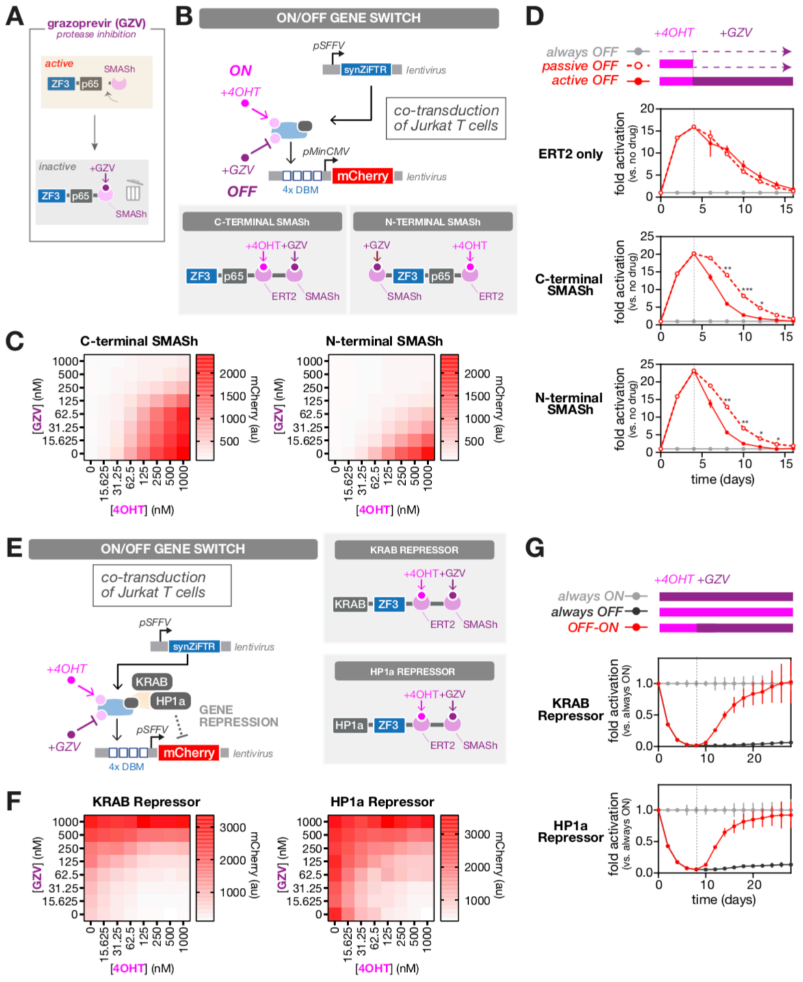


**Fig. S5. Design and characterization of synZiFTR ON/OFF switches**

(**A**) Incorporating a SMASh (Small Molecule-Assisted Shutoff) domain to enable GZV-dependent degradation of synZiFTRs. (**B**) Design of a ON/OFF synZiFTR switch that utilizes both ERT2 and SMASh domains for 4OHT-inducible ON and GZV-inducible OFF gene expression switching. Jurkat T cells were co-transduced with reporter and synZiFTR expression lentiviral vectors in an equal ratio. SMASh domains were fused either C-terminally or N-terminally. (**C**) Two-input dose-response profiles for the ON/OFF switch. Dose-responses were generated by measuring mCherry by flow cytometry 4 days following induction with different combinations of GZV and 4OHT. Fold activation levels represent mean values for three biological replicates. (**D**) ON → OFF switching dynamics. We measured reporter activation and de-activation over time for (1) an ERT2-synZiFTR (ON) switch, (2) ON/OFF switch (C-terminal SMASh), (3) ON/OFF switch (N-terminal SMASh). Cells were induced with 1 uM 4OHT for 4 days, washed, and then cultured either without inducer ("passive OFF") or with 1 uM GZV ("active OFF"). (**E**) Design of an ON/OFF synZiFTR switch that utilizes both ERT2 and SMASh domains for 4OHT-inducible OFF and GZV-inducible ON gene expression switching. This design is based on synZiFTR repressors utilizing KRAB or Hp1a domains. Jurkat T cells were co-transduced with reporter and synZiFTR expression lentiviral vectors in an equal ratio. Response promoters contained a DBM array upstream of a constitutive SFFV promoter. (**F**) Two-input dose-response profiles for the ON/OFF switch. Dose-responses were generated by measuring mCherry by flow cytometry 8 days following induction with different combinations of GZV and 4OHT. Fold activation levels represent mean values for three biological replicates. (**G**) OFF → ON switching dynamics. We measured reporter de-activation and re-activation over time for (1) ON/OFF switch utilizing a KRAB-based synZiFTR repressor, (2) ON/OFF switch utilizing a Hp1a-based synZiFTR repressor. Cells were induced with either 1 uM GZV ("always ON"), 1 uM 4OHT ("always OFF"), or 1 uM 4OHT for 8 days followed by 1 uM GZV ("OFF-ON").

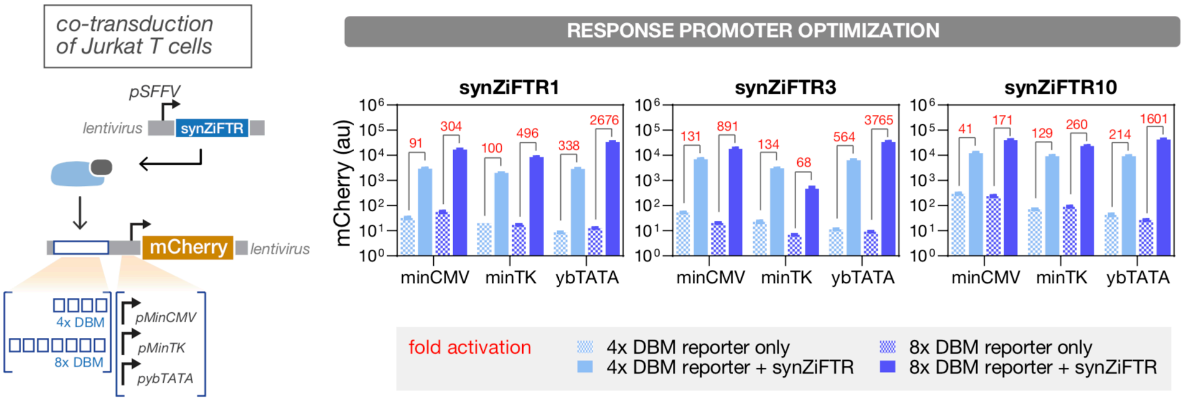


**Fig. S6. Optimization of synZiFTR response promoter architecture**

Schematic of genetic circuit used to screen and optimize response promoters in Jurkat T cells (left). Jurkat T cells were co-transduced with reporter and synZiFTR expression lentiviral vectors in an equal ratio. We screened response promoter configurations composed of 4x or 8x DBM arrays upstream of one of three minimal promoters (minCMV, minTK, and ybTATA). Activation of response promoter configurations by three representative synZiFTRs (right). Bars represent mean values for three measurements ± SD. Fold activation levels between activated and basal conditions are shown in red.

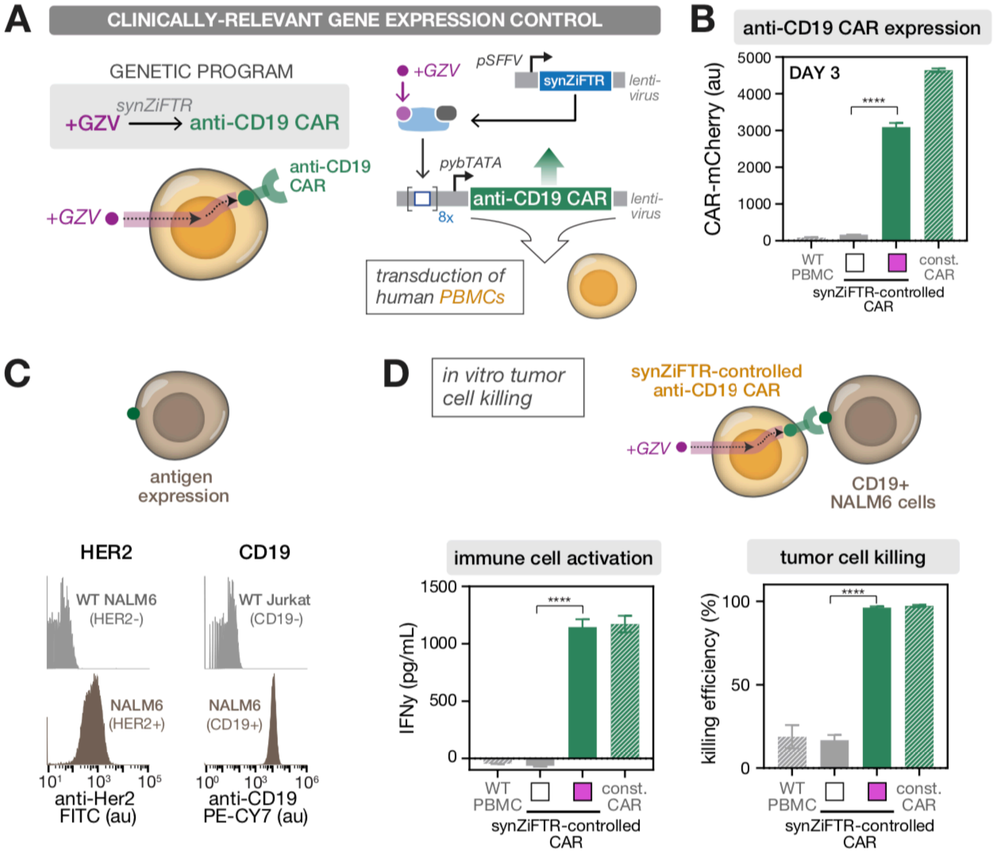


**Fig. S7. SynZiFTRs enable inducer-dependent gene expression control of anti-CD19 CAR T cell activity** (**A**) Conceptualization (left) and implementation (right) of a synZiFTR-controlled anti-CD19 CAR gene expression program. Human PBMCs were activated and co-transduced with equal ratios of lentiviral vectors encoding anti-CD19 CAR payload and synZiFTR expression cassettes. (**B**) SynZiFTR program enables GZV-dependent CAR expression in primary human immune cells. Expression of anti-CD19 CAR-mCherry was measured by flow cytometry two days following induction (with or without 1 uM GZV). Bars represent mean values for three measurements ± SD. Statistics represent two-tailed Student's t test; ***: $p < 0.001$ ; ****: $p < 0.0001$. Const. CAR, constitutively expressed pSFFV-CAR. (**C**) Quantification of surface antigen expression on NALM6 tumor cells. HER2+ and CD19+ NALM6 cells were compared to WT (HER2-) NALM6 and WT (CD19-) Jurkat cells, respectively. (**D**) SynZiFTR-CAR program enables GZV-dependent immune cell activation and tumor cell killing *in vitro*. SynZiFTR-controlled CAR cells (pre-induced with or without 1 uM GZV for 2 days) were co-cultured with CD19+ NALM6 target leukemia cells in a 1:1 ratio (left). IFNγ secretion from activated immune cells was measured by ELISA (center) and tumor cell killing by flow cytometry (right), one day following co-culturing.

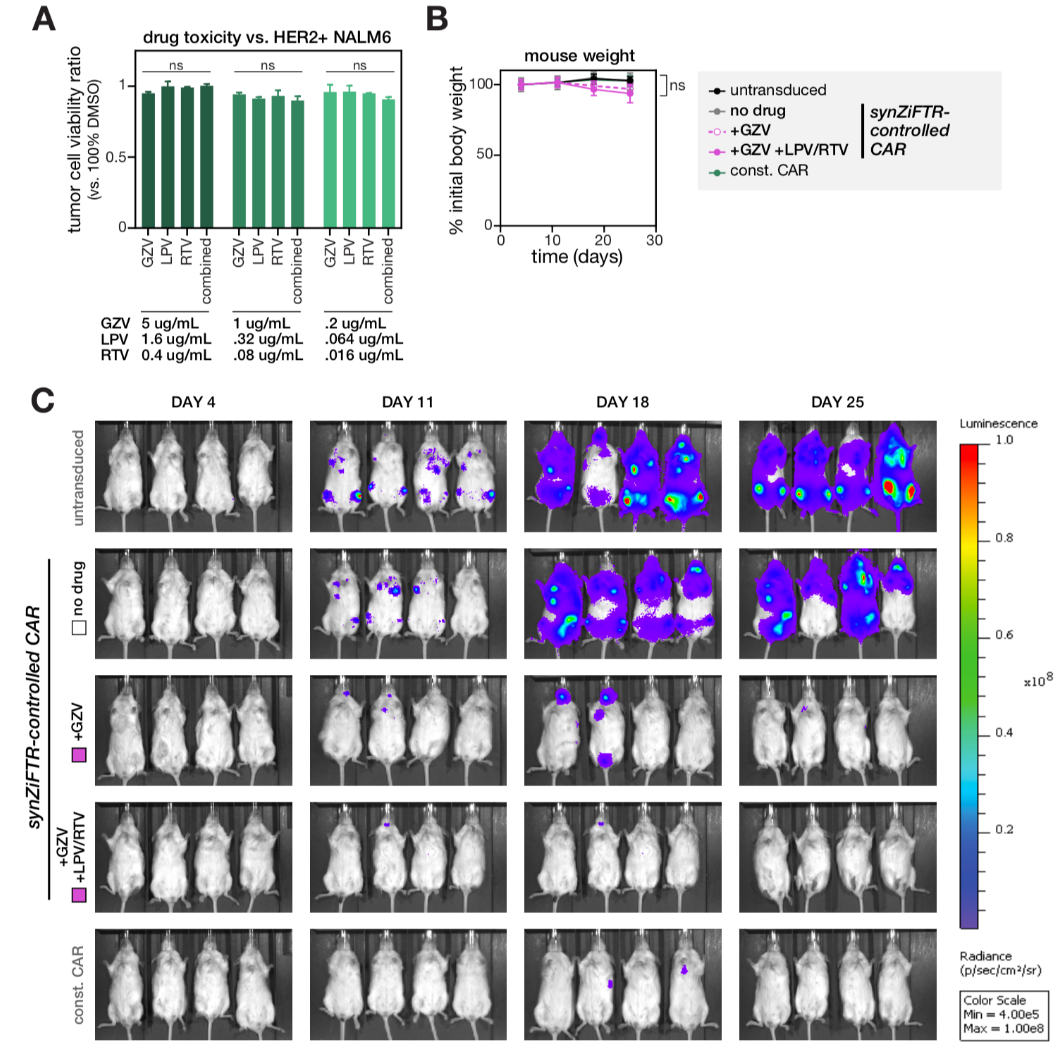


**Fig S8. Testing *in vivo* efficacy of synZiFTR-controlled CAR cells using a xenograft blood tumor mouse model**

(**A**) Small molecule inducer formulations show no cytotoxicity *in vitro*. HER2+ NALM6 cells were cultured with the indicated concentrations of small molecule inducers for 16 hours, and cell viability was quantified by flow cytometry. Bars represent mean values for three measurements ± SD. Statistics represent two-way ANOVA with Tukey's Multiple Comparisons; ns: not significant. (**B**) Change in mouse body weight over time, measured as the percentage of body weight to the initial weight on day 0. Points represent mean values ± SD (n=4 mice per condition). (**C**) IVIS imaging of mouse groups treated with (1) untransduced PBMCs, (2) synZiFTR-controlled CAR cells, (3) synZiFTR-controlled CAR cells with GZV, (4) synZiFTR-controlled CAR cells with GZV+LPV/RTV, (4) constitutive CAR cells. (n=4 mice per condition). Mice were imaged on days 4, 11, 18, and 25 post PBMC injection. Related to **Fig. 3F**.

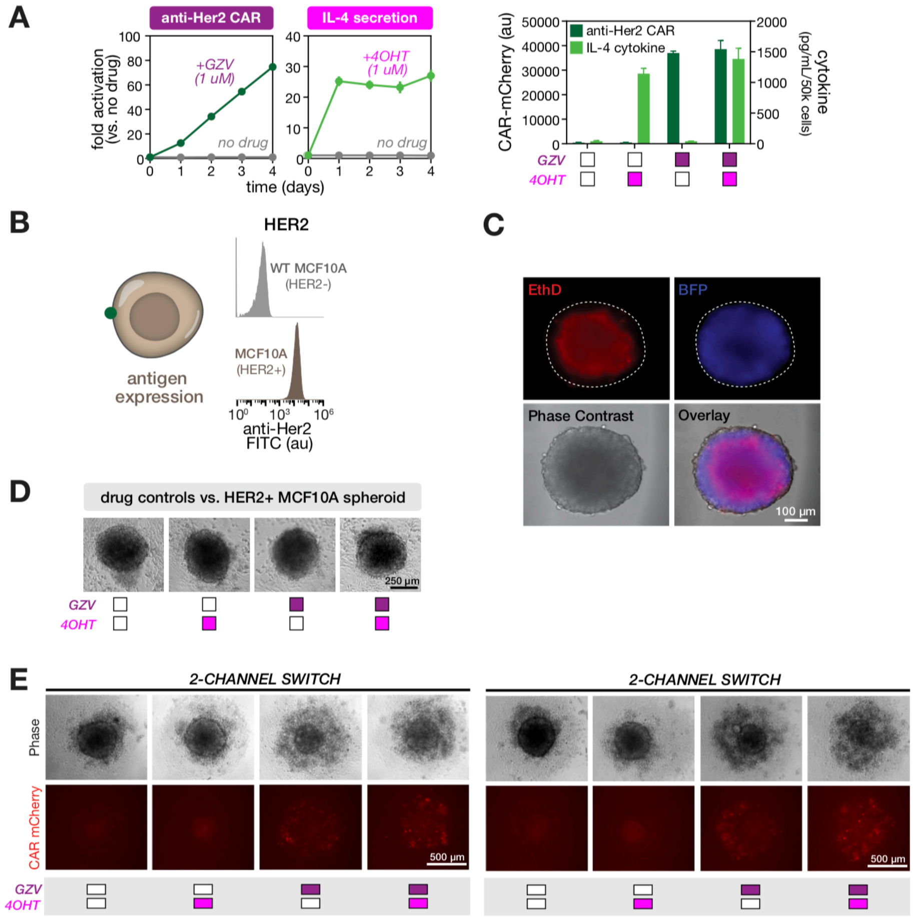

**Fig. S9. Testing the efficacy of two-channel synthetic switch cells in 3D tumor spheroids**
(**A**) Independent, drug-inducible control of anti-Her2 CAR and IL-12 in primary human immune cells. PBMCs were transduced with lentiviral vectors comprising the two-channel system, and induced with combinations of GZV and 4OHT. Anti-Her2-CAR-mCherry was measured by flow cytometry and cytokine secretion by ELISA. Points (left) and bars (right) represent mean values for three measurements ± SD. (**B**) Quantification of surface antigen expression on MCF10A breast mammary epithelial cells. Her2-overexpressing (HER2+) MCF10A cells were compared to WT (HER2-) MCF10A cells. (**C**) HER2+ MCF10A BFP spheroids show the development of a central necrotic core after 5 days of culture. Fluorescence imaging shows a viable, BFP-expressing peripheral layer of cells surrounding a centralized region of dead cells. (**D**) Spheroids show no morphological disruption in the presence of small molecule inducers. 5 day old spheroids were cultured in PBMC media with the indicated combination of inducers. +GZV, 1 uM; +4OHT, 1 uM. (**E**) Biological replicates of spheroid / PBMC co-culture showing GZV-inducible control over spheroid destruction by engineered two-channel cells. Representative phase contrast and CAR-mCherry fluorescent images of spheroid morphology when co-cultured with two-channel inducible or control PBMCs. Clear disruption of the compact, rounded morphology is seen in conditions in which PBMC express the anti-HER2 CAR.

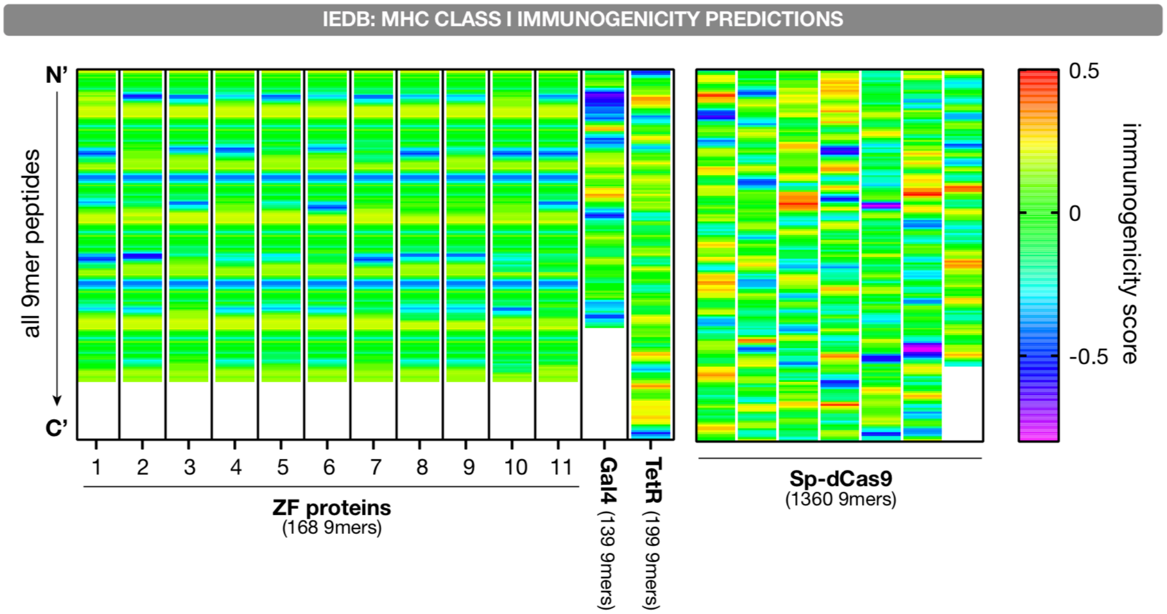


**Fig. S10. Predicted immunogenicity of epitope sequences in engineered ZFs and commonly-used DNA-binding domains**

We used the T Cell Class I pMHC Immunogenicity tool from the Immune Epitope Database (IEDB) to evaluate the immunogenicity scores of all 9-mer peptide sequences within the 11 engineered ZFs and other commonly-used DNA-binding domains (Gal4, TetR, and *S. pyogenes* dCas9). A higher immunogenicity score for a peptide indicates that its composition more closely resembles that of immunogenic peptides, and thus indicates a higher probability of eliciting an immune response. Sp-dCas9, *S. pyogenes* dCas9.

#### Supplementary Tables

##### Table S1. DNA-binding motif sequences

| DNA-Binding Motif (DNA-Binding Protein) | DNA Sequence (5' => 3') | Reference |
|---|---|---|
| DBM 1 (ZF1) | cGTCGAAGTCGAAGTCGACc | This study |
| DBM 2 (ZF2) | gGACGACGTTACGGACGTAc | This study |
| DBM 3 (ZF3) | aGACGTCGAAGTAGCCGTAg | This study |
| DBM 4 (ZF4) | gGACGACGCCGATGTAGAAg | This study |
| DBM 5 (ZF5) | tGAAGCAGTCGACGCCGAAg | This study |
| DBM 6 (ZF6) | gGACGACGCGGTCTAAGAAg | This study |
| DBM 7 (ZF7) | cGACGAGGTCGCATAAGTAg | This study |
| DBM 8 (ZF8) | aGACGCAGTATAGGTCGAAc | This study |
| DBM 9 (ZF9) | aGACGCAGTATAGGACGACg | This study |
| DBM 10 (ZF10) | cGGCGTAGCCGATGTCGCGc | This study |
| DBM 11 (ZF11) | gGTCGTTGCGGTAGTCGAAg | This study |
| UAS (Gal4) | GGAGCACTGTCCTCCGAACG | Morsut et al., 2016 |
| TetO (TetR) | TCCCTATCAGTGATAGAGA | Morsut et al., 2016 |
| ZFHD1 BS (ZFHD1) | TAATGATGGGCG | Morsut et al., 2016 |

#### Table S2. Protein sequences of engineered ZFs and other DNA-binding domains

| DNA-Binding Domain | Protein Sequence (N' => C') | Reference |
|---|---|---|
| ZF1 (ZF1-3) | SRPGERPFQCRICMRNFSEEANLRRHTRTHTGEKPFQCRICMRNFSDHSSLKRHLRTHTGSQKPFQCRICMRNFSQSANLLRHTRTHTGEKPFQCRICMRNFSDPSSLKRHLRTHTGSQKPFQCRICMRNFSQQTNLTRHTRTHTGEKPFQCRICMRNFSDATQLVRHLRTHLRGS | This study |
| ZF2 (ZF2-6) | SRPGERPFQCRICMRNFSQKQALTRHTRTHTGEKPFQCRICMRNFSDRGNLTRHLRTHTGSQKPFQCRICMRNFSRSHDLTRHTRTHTGEKPFQCRICMRNFSHKSSLTRHLRTHTGSQKPFQCRICMRNFSEKQNLARHTRTHTGEKPFQCRICMRNFSDPSNLRRHLRTHLRGS | This study |
| ZF3 (ZF3-5) | SRPGERPFQCRICMRNFSQRSSLVRHTRTHTGEKPFQCRICMRNFSDKSVLARHLRTHTGSQKPFQCRICMRNFSQRSSLVRHTRTHTGEKPFQCRICMRNFSQRNNLGRHLRTHTGSQKPFQCRICMRNFSTHAVLTRHTRTHTGEKPFQCRICMRNFSDRGNLTRHLRTHLRGS | This study |
| ZF4 (ZF4-8) | SRPGERPFQCRICMRNFSQQTNLTRHTRTHTGEKPFQCRICMRNFSQSTSLQRHLRTHTGSQKPFQCRICMRNFSVRHNLTRHTRTHTGEKPFQCRICMRNFSDHSNLSRHLRTHTGSQKPFQCRICMRNFSEKQNLARHTRTHTGEKPFQCRICMRNFSDPSNLRRHLRTHLRGS | This study |
| ZF5 (ZF5-7) | SRPGERPFQCRICMRNFSNMSNLTRHTRTHTGEKPFQCRICMRNFSDRSVLRRHLRTHTGSQKPFQCRICMRNFSDPSNLARHTRTHTGEKPFQCRICMRNFSDRSSLRRHLRTHTGSQKPFQCRICMRNFSQSGTLHRHTRTHTGEKPFQCRICMRNFSQRPNLTRHLRTHLRGS | This study |
| ZF6 (ZF6-4) | SRPGERPFQCRICMRNFSQQTNLTRHTRTHTGEKPFQCRICMRNFSQKANLGVHLRTHTGSQKPFQCRICMRNFSDHSSLKRHTRTHTGEKPFQCRICMRNFSRADMLRRHLRTHTGSQKPFQCRICMRNFSEKQNLARHTRTHTGEKPFQCRICMRNFSDPSNLRRHLRTHLRGS | This study |
| ZF7 (ZF7-3) | SRPGERPFQCRICMRNFSQRSSLVRHTRTHTGEKPFQCRICMRNFSQRGNLNMHLRTHTGSQKPFQCRICMRNFSRRQELTRHTRTHTGEKPFQCRICMRNFSDHSSLKRHLRTHTGSQKPFQCRICMRNFSRQDNLGRHTRTHTGEKPFQCRICMRNFSDGGNLGRHLRTHLRGS | This study |
| ZF8 (ZF8-1) | SRPGERPFQCRICMRNFSQASNLTRHTRTHTGEKPFQCRICMRNFSDHSSLKRHLRTHTGSQKPFQCRICMRNFSRAHNLLLHTRTHTGEKPFQCRICMRNFSQRSSLVRHLRTHTGSQKPFQCRICMRNFSQSTTLKRHTRTHTGEKPFQCRICMRNFSDPSNLRRHLRTHLRGS | This study |
| ZF9 (ZF9-2) | SRPGERPFQCRICMRNFSEKQNLARHTRTHTGEKPFQCRICMRNFSDPSNLRRHLRTHTGSQKPFQCRICMRNFSRAHNLLLHTRTHTGEKPFQCRICMRNFSQRSSLVRHLRTHTGSQKPFQCRICMRNFSQSTTLKRHTRTHTGEKPFQCRICMRNFSDPSNLRRHLRTHLRGS | This study |
| ZF10 (ZF10-1) | SRPGERPFQCRICMRNFSRRHGLDRHTRTHTGEKPFQCRICMRNFSDHSSLKRHLRTHTGSQKPFQCRICMRNFSVRHNLTRHLRTHTGEKPFQCRICMRNFSDHSNLSRHLKTHTGSQKPFQCRICMR | This study |

| | | |
|---|---|---|
| | NFSQRSSLVRHLRTHTGEKPFQCRICMRNFSESGHLKRHLRTHLRGS | |
| ZF11 (ZF11-1) | SRPGERPFQCRICMRNFSQLSNLTRHTRTHTGEKPFQCRICMRNFSDRSSLKRHLRTHTGSQKPFQCRICMRNFSQRSSLVRHLRTHTGEKPFQCRICMRNFSRLDMLARHLKTHTGSQKPFQCRICMRNFSVRHSLTRHLRTHTGEKPFQCRICMRNFSESGALRRHLRTHLRGS | This study |
| GAL4 DBD | MKLLSSIEQACDICRLKKLKCSKEKPKCAKCLKNNWECRYSPKTKRSPLTRAHLTEVESRLERLEQLFLLIFPREDLDMILKMDSLQDIKALLTGLFVQDNVNKDAVTDRLASVETDMPLTLRQHRISATSSSEESSNKGQRQLTVS | Morsut et al., 2016 |
| TetR DBD | MSRLDKSKVINSALELLNEVGIEGLTTRKLAQKLGVEQPTLYWHVKNKRALLDALAIEMLDRHHTHFCPLEGESWQDFLRNNAKSFRCALLSHRDGAKVHLGTRPTEKQYETLENQLAFLCQQGFSLENALYALSAVGHFTLGCVLEDQEHQVAKEERETPTTDSMPPLLRQAIELFDHQGAEPAFLFGLELIICGLEKQLKCESGG | Morsut et al., 2016 |
| Sp-dCas9 DBD | MDKKYSIGLAIGTNSVGWAVITDEYKVPSKKFKVLGNTDRHSIKKNLIGALLFDSGETAEATRLKRTARRRYTRRKNRICYLQEIFSNEMAKVDDSFFHRLEESFLVEEDKKHERHPIFGNIVDEVAYHEKYPTIYHLRKKLVDSTDKADLRLIYLALAHMIKFRGHFLIEGDLNPDNSDVDKLFIQLVQTYNQLFEENPINASGVDAKAILSARLSKSRRLENLIAQLPGEKKNGLFGNLIALSLGLTPNFKSNFDLAEDAKLQLSKDTYDDDLDNLLAQIGDQYADLFLAAKNLSDAILLSDILRVNTEITKAPLSASMIKRYDEHHQDLTLLKALVRQQLPEKYKEIFFDQSKNGYAGYIDGGASQEEFYKFIKPILEKMDGTEELLVKLNREDLLRKQRTFDNGSIPHQIHLGELHAILRRQEDFYPFLKDNREKIEKILTFRIPYYVGPLARGNSRFAWMTRKSEETITPWNFEEVVDKGASAQSFIERMTNFDKNLPNEKVLPKHSLLYEYFTVYNELTKVKYVTEGMRKPAFLSGEQKKAIVDLLFKTNRKVTVKQLKEDYFKKIECFDSVEISGVEDRFNASLGTYHDLLKIIKDKDFLDNEENEDILEDIVLTLTLFEDREMIEERLKTYAHLFDDKVMKQLKRRRYTGWGRLSRKLINGIRDKQSGKTILDFLKSDGFANRNFMQLIHDDSLTFKEDIQKAQVSGQGDSLHEHIANLAGSPAIKKGILQTVKVVDELVKVMGRHKPENIVIEMARENQTTQKGQKNSRERMKRIEEGIKELGSQILKEHPVENTQLQNEKLYLYYLQNGRDMYVDQELDINRLSDYDVAAIVPQSFLKDDSIDNKVLTRSDKARGKSDNVPSEEVVKKMKNYWRQLLNAKLITQRKFDNLTKAERGGLSELDKAGFIKRQLVETRQITKHVAQILDSRMNTKYDENDKLIREVKVITLKSKLVSDFRKDFQFYKVREINNYHHAHDAYLNAVVGTALIKKYPKLESEFVYGDYKVYDVRKMIAKSEQEIGKATAKYFFYSNIMNFFKTEITLANGEIRKRPLIETNGETGEIVWDKGRDFATVRKVLSMPQVNIVKKTEVQTGGFSKESILPKRNSDKLIARKKDWDPKKYGGFDSPTVAYSVLVVAKVEKGKSKKLKSVKELLGITIMERSSFEKNPIDFLEAKGYKEVKKDLIIKLPKYSLFELENGRKRMLASAGELQKGNELALPSKYVNFLYLASHYEKLKGSPEDNEQKQLFVEQHKHYLDEIIEQISEFSKRVILADANLDKVLSAYNKHRDKPIREQAENIIHLFTLTNLGAPAAFKYFDTTIDRKRYTSTKEVLDATLIHQSITGLYETRIDLSQLGGD | Mali et al., 2013 |

**Table S3. Plasmids used in this study**

See attached Excel spreadsheet

#### References and Notes


1. M. L. Maeder, S. Thibodeau-Beganny, J. D. Sander, D. F. Voytas, J. K. Joung, Oligomerized pool engineering (OPEN): an 'open-source' protocol for making customized zinc-finger arrays. *Nat Protoc* **4**, 1471-1501 (2009).
2. J. D. Sander *et al.*, Selection-free zinc-finger-nuclease engineering by context-dependent assembly (CoDA). *Nat Methods* **8**, 67-69 (2011).
3. S. Thibodeau-Beganny, J. K. Joung, Engineering Cys2His2 zinc finger domains using a bacterial cell-based two-hybrid selection system. *Methods Mol Biol* **408**, 317-334 (2007).
4. M. Isalan, Y. Choo, A. Klug, Synergy between adjacent zinc fingers in sequence-specific DNA recognition. *Proc Natl Acad Sci U S A* **94**, 5617-5621 (1997).
5. M. Isalan, A. Klug, Y. Choo, Comprehensive DNA recognition through concerted interactions from adjacent zinc fingers. *Biochemistry* **37**, 12026-12033 (1998).
6. J. S. Kim, C. O. Pabo, Getting a handhold on DNA: design of poly-zinc finger proteins with femtomolar dissociation constants. *Proc Natl Acad Sci U S A* **95**, 2812-2817 (1998).
7. M. Moore, A. Klug, Y. Choo, Improved DNA binding specificity from polyzinc finger peptides by using strings of two-finger units. *Proc Natl Acad Sci U S A* **98**, 1437-1441 (2001).
8. D. E. Paschon *et al.*, Diversifying the structure of zinc finger nucleases for high-precision genome editing. *Nat Commun* **10**, 1133 (2019).
9. D. J. Segal *et al.*, Evaluation of a modular strategy for the construction of novel polydactyl zinc finger DNA-binding proteins. *Biochemistry* **42**, 2137-2148 (2003).
10. H. A. Greisman, C. O. Pabo, A general strategy for selecting high-affinity zinc finger proteins for diverse DNA target sites. *Science* **275**, 657-661 (1997).
11. N. P. Pavletich, C. O. Pabo, Zinc finger-DNA recognition: crystal structure of a Zif268-DNA complex at 2.1 A. *Science* **252**, 809-817 (1991).
12. L. Morsut *et al.*, Engineering Customized Cell Sensing and Response Behaviors Using Synthetic Notch Receptors. *Cell* **164**, 780-791 (2016).
13. J. J. Calis *et al.*, Properties of MHC class I presented peptides that enhance immunogenicity. *PLoS computational biology* **9**, e1003266 (2013).
14. M. Park, N. Patel, A. J. Keung, A. S. Khalil, Engineering Epigenetic Regulation Using Synthetic Read-Write Modules. *Cell* **176**, 227-238 e220 (2019).
15. R. Zufferey *et al.*, Self-inactivating lentivirus vector for safe and efficient in vivo gene delivery. *Journal of virology* **72**, 9873-9880 (1998).
16. A. Casini *et al.*, R2oDNA designer: computational design of biologically neutral synthetic DNA sequences. *ACS Synth Biol* **3**, 525-528 (2014).
17. W. J. Kent *et al.*, The human genome browser at UCSC. *Genome Res* **12**, 996-1006 (2002).
18. B. Langmead, S. L. Salzberg, Fast gapped-read alignment with Bowtie 2. *Nat Methods* **9**, 357-359 (2012).
19. B. Langmead, C. Trapnell, M. Pop, S. L. Salzberg, Ultrafast and memory-efficient alignment of short DNA sequences to the human genome. *Genome Biol* **10**, R25 (2009).
20. M. I. Love, W. Huber, S. Anders, Moderated estimation of fold change and dispersion for RNA-seq data with DESeq2. *Genome Biol* **15**, 550 (2014).
21. T. Tanaka, H. Nakazawa, N. Kuriyama, M. Kaneki, Farnesyltransferase inhibitors prevent HIV protease inhibitor (lopinavir/ritonavir)-induced lipodystrophy and metabolic syndrome in mice. *Exp Ther Med* **15**, 1314-1320 (2018).
